# Causal mediation analysis for time-varying heritable risk factors with Mendelian Randomization

**DOI:** 10.1101/2024.02.10.579129

**Authors:** Zixuan Wu, Ethan Lewis, Qingyuan Zhao, Jingshu Wang

## Abstract

Understanding the causal mechanisms of diseases is crucial in clinical research. When randomized experiments are unavailable, Mendelian Randomization (MR) leverages genetic mutations to mitigate confounding. However, most MR analyses assume static risk factors, oversimplifying dynamic risk factor effects. The framework of life-course MR addresses this but struggles with limited GWAS cohort sizes and correlations across time points. We propose FLOW-MR, a computational approach estimating causal structural equations for temporally ordered traits using only GWAS summary statistics. FLOW-MR enables inference on direct, indirect, and path-wise causal effects, demonstrating superior efficiency and reliability, especially with noisy data. By incorporating a spike-and-slab prior, it mitigates challenges from extreme polygenicity and weak instruments. Applying FLOW-MR, we uncovered a childhood-specific protective effect of BMI on breast cancer and analyzed the evolving impacts of BMI, systolic blood pressure, and cholesterol on stroke risk, revealing their causal relationships.

## Introduction

Understanding the causal mechanisms behind diseases is a core challenge in clinical research. While randomized controlled trials are considered the gold standard for establishing causality, they can be impractical in certain situations. As a result, Mendelian Randomization (MR) has become increasingly popular as an alternative. MR leverages genetic variations as a form of “natural experiment”, effectively reducing the influence of unmeasured environmental confounders in epidemiological studies [1, 2].

Traditional MR analyses often rely on cross-sectional designs, treating risk factors as constant over time. However, many heritable risk factors vary across the lifespan and may have time-specific effects on disease outcomes [3, 4]. Ignoring this temporal aspect can lead to simplistic and misleading conclusions [5]. For example, while ecological studies in epidemiology show that vitamin D levels during childhood are associated with multiple sclerosis risk, standard MR approaches, which focus on adult vitamin D levels, have instead pointed to adult vitamin D as the key factor in disease etiology [6].

In response to these limitations, life-course MR has emerged as a framework that considers how risk factors measured across an individual’s lifetime influence later-life outcomes [7]. Though the idea of incorporating life-long information is appealing, performing life-course MR can be challenging due to the small cohort size in Genome-wide Association Studies (GWAS) of earlier life traits and high auto-correlations of risk factors over time. One commonly used approach is multivariable MR (MVMR), such as MVMR-IVW [8], which estimates the *direct* causal effects of the risk factor at each time point. However, MVMR will have limited efficiency and reliability on noisy GWAS data with a small cohort size. Additionally, current MVMR techniques struggle to evaluate indirect causal effects of early-life risk factors that influence outcomes through intermediate factors. Other methods, such as g-estimations of structural mean models [4, 9] or functional principal component analysis that aggregates effects across time points [10], offer some alternative solutions but require access to individual-level GWAS data that are often not available on a large scale.

To address these challenges, we propose FLOW-MR (Framework for Life-cOurse pathWay analysis using Mendelian Randomization), a computational method that estimates a full system of structural equations for temporally ordered heritable traits using only GWAS summary statistics. FLOW-MR enables comprehensive causal mediation analysis, allowing for the decomposition of total causal effects into direct, indirect, and path-specific components. As a longitudinal extension of GRAPPLE [11], FLOW-MR introduces a novel sufficient condition for causal identifiability that generalizes the well-known InSIDE assumption [12] to longitudinal settings, thereby accommodating both pervasive horizontal pleiotropy and unmeasured mediator-outcome confounding. FLOW-MR can be viewed as a Bayesian analogue of the full-information maximum likelihood (FIML) estimator commonly used in structural equation modeling in economics and social sciences [13], with the advantage of requiring only summary data. Its hierarchical Bayesian framework allows for flexible modeling of weak and invalid instruments and captures complex interactions between genetic variants and environmental factors.

When benchmarked against existing MVMR methods, FLOW-MR substantially improves the efficiency and robustness of the statistical inference, especially when the GWAS data are noisy and from smaller cohorts. By incorporating a spike-and-slab prior for genetic effects, FLOW-MR can tackle the different polygenic patterns of complex traits and mitigate weak instrument bias. Applying FLOW-MR to publicly available GWAS summary statistics, we identify a protective effect of Body Mass Index (BMI) during childhood on breast cancer risk, confined to a specific developmental period. Our analysis further reveals a complex interplay between BMI, systolic blood pressure (SBP), and low-density cholesterol levels across different life stages and their effects on stroke risk. Our findings suggest that adulthood SBP may be the only factor with a direct causal effect on stroke, while the previously identified causal effect of BMI on SBP [14, 15] in adulthood may be attributable to confounding by childhood SBP.

## Results

### Method overview

Consider a sequence of risk factors (*X*_1_, *X*_2_, …, *X*_*K*−1_) in temporal order, so that *X*_*i*_ precedes *X*_*j*_ when *i < j*. Our goal is to understand the causal relationship between these traits and their effects on a later outcome *Y* (also denoted as *X*_*K*_ for convenience), typically some disease status measured in adulthood. Causal effects among the traits must adhere to the temporal sequence (i.e., only an earlier trait can causally influence a later trait) (Figure 1a). We allow the genetic variants ***Z*** and the unmeasured environmental confounders ***U*** to have direct causal effects on any trait at any time point. Associated with this directed acyclic graph (DAG), we assume a series of structural equations for each individual (see Methods).

**Figure 1:**
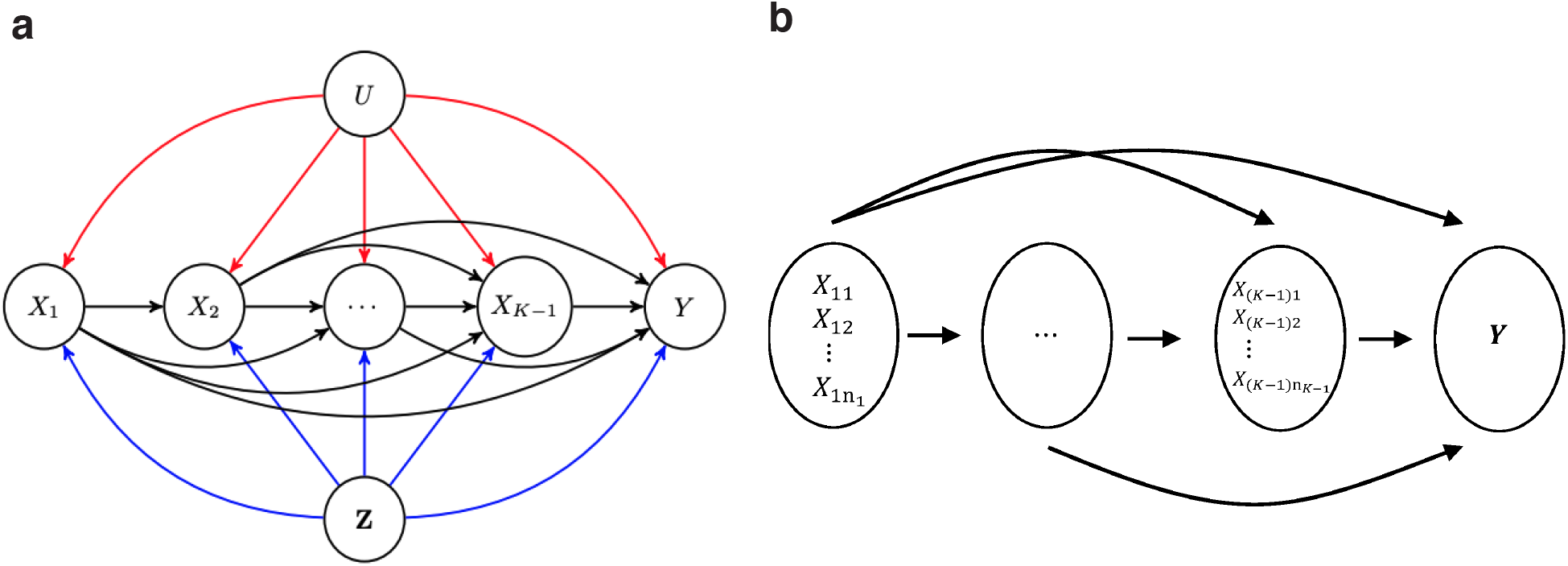
Model overview. a) *X*_1_, …, *X*_*K*−1_ are exposure traits in increasing temporal order and *Y* is the outcome trait. The blue arrows indicate the causal effects of the genetic variants ***Z*** on the traits. The red arrows represent the effects of unmeasured non-heritable confounders ***U***. b) Illustration of causal relationships across traits, allowing for multiple traits at each time point.

FLOW-MR requires only GWAS summary statistics for each trait, achieved by projecting the individual-level structural equations onto each single SNP *Z*_*j*_. Specifically, let *γ*_*kj*_ be the marginal associations between a SNP *j* and trait *X*_*k*_, we then have the following linear models:

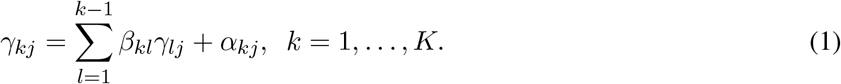

Here, *β*_*kl*_ is our parameter of interest, representing the direct causal effect of an earlier trait *X*_*l*_ on a later trait *X*_*k*_. The term *α*_*kj*_ denotes the “direct” genetic effect of SNP *j* on trait *k*. If SNP *Z*_*j*_ is used as an instrument for trait *X*_*k*_, then *α*_*kj*_ corresponds to the SNP-trait link used for instrumental variable analysis. In contrast, for 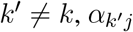 represents pleiotropic effects of SNP *Z*_*j*_ on trait *X*_*k*′_.

GWAS summary statistics provide estimates of the marginal associations *γ*_*kj*_ with their standard errors. They can be obtained either from linear regressions or, for binary traits, from logistic regressions, following [16] under the liability threshold model (Figure S1, SI text). For each SNP *Z*_*j*_ and trait *X*_*k*_, we assume 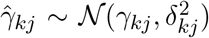, where 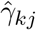 is the observed estimate and 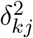 is the estimated variance. Our approach accommodates sample overlap across traits, thus allowing for correlations among 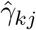 across different traits *k*. Following previous MR methods [11, 17], we can adaptively estimate this correlation matrix from the GWAS summary data.

Notice that, unlike marginal associations *γ*_*kj*_, the “direct” genetic effects *α*_*kj*_ are not observed from the data. Existing literature shows that most SNPs may have non-zero but weak effects on complex traits, while a small subset may be responsible for the core biological process and strongly impact the traits [11]. Therefore, FLOW-MR assumes the following spike-and-slab distributional assumptions on *α*_*kj*_:

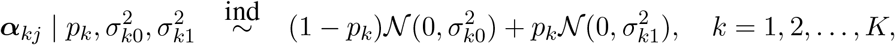

which allows for trait-specific effect sizes. To estimate the model, we use a hierarchical Bayesian framework and Gibbs sampler. The Gibbs sampler enables the convenient use of posterior samples to construct credible intervals for any function of the direct effects {*β*_*kl*_}_*k*≥*l*_, including both direct and indirect effects among the traits, as well as the proportions of these effects relative to the total effects.

To address known confounders, we further extend our model to accommodate multiple traits at any given time point (Figure 1b). Specifically, at each time point *k*, we assume that there are *n*_*k*_ ≥ 1 exposures, denoted as 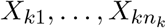. To facilitate the MR analysis, we assume that no causal relationship exists between traits measured at the same time point. If a known causal direction exists between two traits measured at the same time point *k*, our method can be still be applied by introducing a “pseudo-time point” (or stage) *k* + 1 following *k*, where the outcome trait of the two traits at time *k* is moved to stage *k* + 1. More details on the model, estimation, inference procedures, and key assumptions are provided in Methods and SI Text.

### Identifiability of direct and indirect causal effects

Mediation analysis, as conducted in this study, is inherently vulnerable to unmeasured mediator-outcome confounding even when the treatment is fully randomized [18]. As illustrated in Figure 2, with two exposures and an outcome (*K* = 3), if we rely solely on a set of valid genetic instruments for the first risk factor *X*_1_ (Figure 2a), it is not possible to separate the direct and indirect effects of *X*_1_ due to unmeasured mediator-outcome confounding between *X*_2_ and *Y* (see Supplementary Text for a counter-example). In contrast, if each risk factor has its own set of valid instruments (Figure 2b), it becomes feasible to identify all direct and indirect causal effects. However, since the sequence of risk factors often represents the same factor measured at different time points, finding SNPs that exert their effects exclusively at a specific time point can be challenging.

FLOW-MR requires a more realistic assumption to account for pleiotropic effects of the SNPs (Figure 2c). Instead of assigning each risk factor its own set of valid instruments, we allow SNPs to have nonzero direct effects on all risk factors—thereby permitting any given SNP to be an invalid instrument for a particular trait. However, we require that the “direct” genetic effects 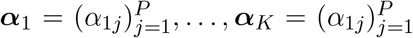 be orthogonal across traits (see Methods). More generally, if the number of SNPs *P* is large, it suffices for these direct effects to be independent rather than exactly orthogonal, extending the well-known InSIDE assumption [12] to multiple traits. Notice that we only assume independence among *α*_*kj*_, allowing the marginal associations *γ*_*kj*_ to remain possibly correlated across traits. This assumption enables FLOW-MR to address both pleiotropic effects and unmeasured mediator-outcome confounding. A formal mathematical statement of the identification conditions and its proof are provided in the SI Text.

**Figure 2:**
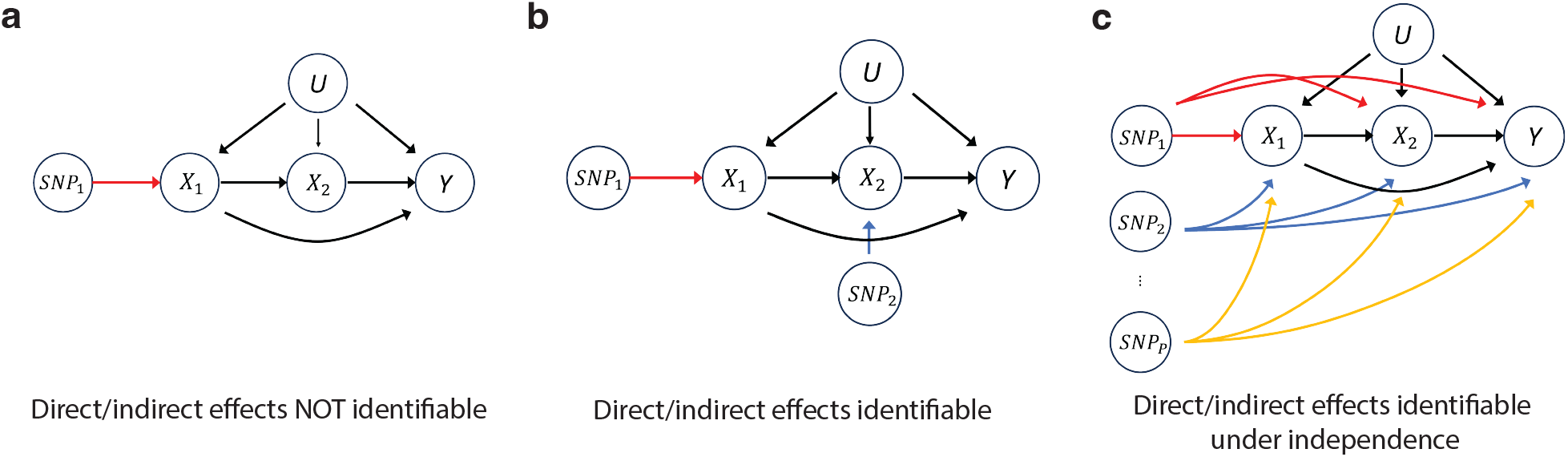
Model identifiability under three scenarios

### Systematic benchmarking with synthetic GWAS summary statistics

We compare FLOW-MR with alternative MVMR approaches to perform life-course MR. Specifically, we benchmark our method against six methods, including MVMR-IVW [8], MVMR-Egger [19], MVMR-Robust [20], GRAPPLE [11], MVMR-Horse [21] and MVMR-cML [22]. Among these methods, MVMR-IVW is the most widely used in practice, and other methods are designed to deal with pleiotropic effects under different assumptions and weak instruments. To apply an MVMR method, we use a multiple-step approach where at each step *k*, we estimate the direct causal effects of *X*_1_, …, *X*_*k*−1_ on *X*_*k*_. One limitation of this approach is the challenge of providing reliable statistical inference on indirect and pathwise effects, as estimates across different steps are correlated.

To assess performance, we generate synthetic GWAS summary datasets by soft-thresholding real GWAS summary statistics, allowing the “true” direct genetic effects to be exactly 0 for some SNPs. Specifically, if we observe 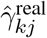 with variance 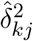 for a particular SNP *j* in a real GWAS dataset, we set 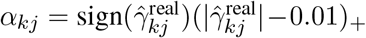 and randomly shuffle *α*_*kj*_ across *j* within each trait *k* to ensure independence across traits.

To specify the causal relationship across the temporally ordered traits, we simulate three scenarios: *K* = 3, *K* = 4, and a multivariate case with multiple exposures at each time point (Figure 3a). We generate synthetic data for earlier traits using childhood GWAS data to mimic real MR mediation analysis, where earlier childhood traits typically have smaller sample sizes than adulthood traits. Specifically, we use childhood BMI data from the Norwegian Mother, Father, and Child Cohort Study (MoBa) [23] and childhood lipid traits from the Avon Longitudinal Study of Parents and Children (ALSPAC) cohort [24]. Adult GWAS datasets are used to generate summary statistics for the remaining traits. Further details of the simulation design, along with additional sensitivity analyses and evaluation of computation costs, can be found in the SI Text. Tables S1-S2 present the number of SNPs and conditional F-statistics [25] computed at each p-value threshold as evaluations of instrument strength.

We evaluate the coverage and average lengths of the confidence and credible intervals of the direct and indirect causal effects across different methods (Figure 3bc, Figures S2-S3). For the direct causal effects, most MVMR methods suffer from severe under-coverage, largely because of limited capabilities to handle pervasive pleiotropic effects and weak-instrument bias. Specifically, MVMR-IVW does not allow any pleiotropic effects, whereas MVMR-Robust and MVMR-cML assume a plurality of valid genetic instruments for each step—a condition that is violated in our simulations and is also unlikely to hold in real life-course MR analyses. Although MVMR-Horse and MVMR-Egger can accommodate independent and pervasive pleiotropy, they remain under-covered in most scenarios, likely due to biases introduced by (conditionally) weak instruments (Table S2) and correlation of SNP-trait associations across risk factors. By contrast, both FLOW-MR and GRAPPLE demonstrate good coverage regardless of SNP strength. Compared to GRAPPLE, FLOW-MR has shorter intervals and offers more efficient analyses. Furthermore, for the indirect causal effects of *X*_1_, only FLOW-MR can provide reliable inference, with credible intervals showing good coverage and reasonable power.

### Effect of early life body size on breast cancer

Recent studies suggest that early-life body size may protect against breast cancer [26–28]. Using MVMR-IVW, [3] found that this protective effect is not mediated by adult body size, which itself does not causally influence breast cancer. However, their study relied on a recalled early-life body score from the UK Biobank as a proxy for early-life body size. As noted by the original authors, such coarse measurements can raise concerns about the accuracy of their findings.

To address this, we revisited the analysis using FLOW-MR and childhood BMI GWAS summary statistics. First, we replicate the original study [3] using the same dataset: adult BMI from the UK Biobank for adult body size, recalled early-life body score from UK Biobank for early-life body size, and breast cancer data from Michailidou et al. [29] for the outcome. FLOW-MR, alongside all MVMR methods, successfully replicates the original findings (Figure 4a, Figure S4a), confirming no causal effect of adult BMI and a direct protective effect of childhood body size on breast cancer.

However, when we replaced the early-life body size trait with GWAS data on 8-year-old BMI, a direct measurement of childhood body size, MVMR-IVW and MVMR-Egger unexpectedly suggested a protective effect of adult BMI on breast cancer, while childhood BMI showed no direct effect (Figure 4b, Figure S4b). In contrast, GRAPPLE, MVMR-Robust and MVMR-Horse lost the power to detect any causal effects. Only FLOW-MR replicates the original conclusion that childhood BMI has a protective effect. Although 8-year-old BMI provides a more precise measurement, its use in MR is challenging due to its small sample size (3K samples in the MoBa cohort compared to half a million in the UK Biobank). This is reflected in the substantial decrease in the conditional F-statistics in the childhood dataset (Table S2), indicating a loss of instrument strength due to the limited cohort size of the MoBa study. FLOW-MR demonstrates both efficiency and robustness in this scenario.

**Figure 3:**
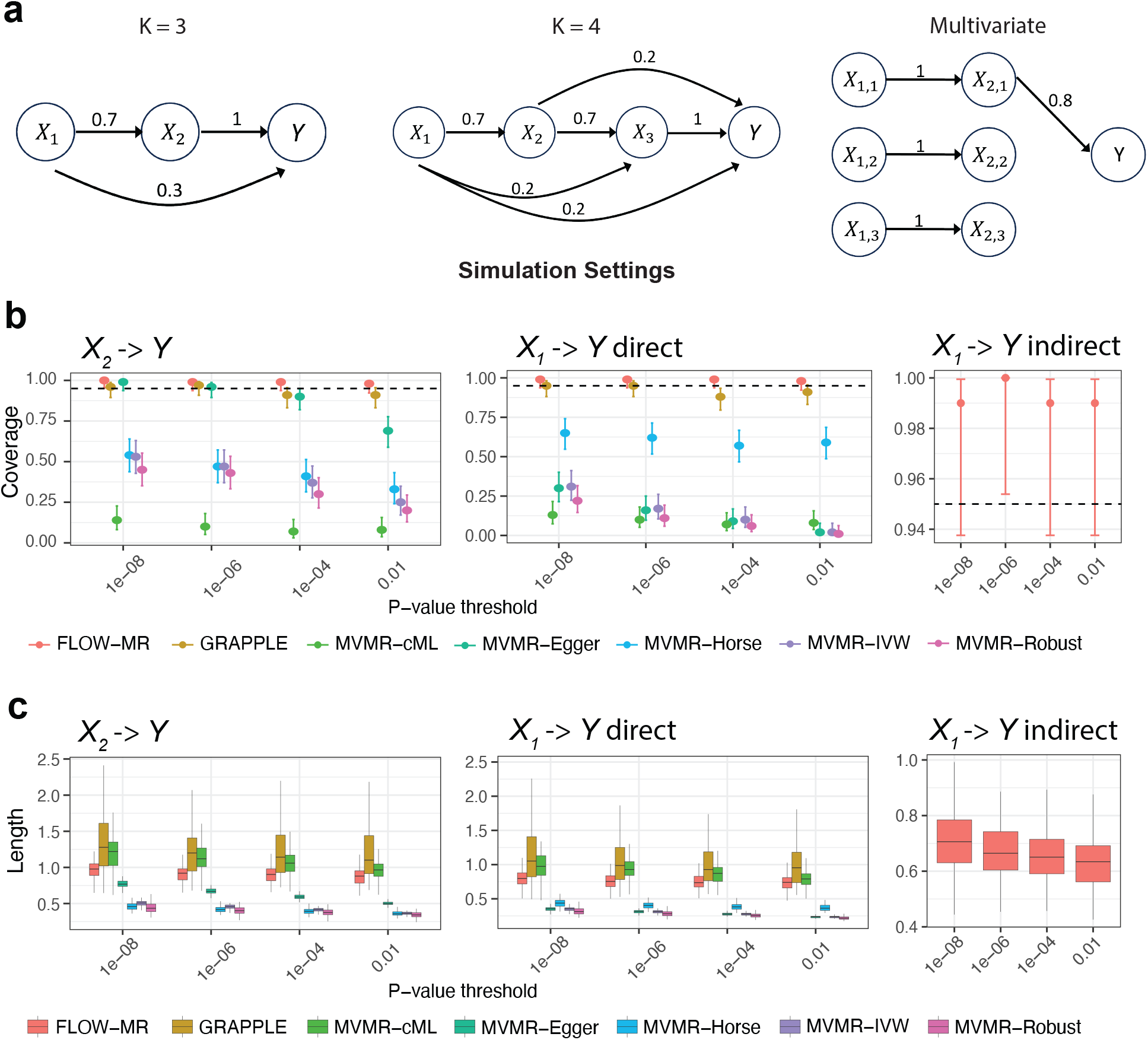
Systematic benchmarking. a) Three simulation scenarios for *K* = 3, *K* = 4 and the multivariate case. b) Empirical coverage of 95% credible intervals on direct effects of *X*_1_ and *X*_2_ on *Y* and indirect effects of *X*_1_ over 100 repeated simulations when *K* = 3. The error bars are the 95% confidence intervals of the coverage. c) Boxplots of lengths of credible intervals over repeated simulations when *K* = 3. Each box represents the lower and upper quartiles, with the line inside indicating the median. The whiskers extend to 1.5 times the interquartile range beyond the box. Outliers have been removed for clarity.

We further explored the impact of including 1-year-old BMI in our analysis. Given that the GWAS data of 1-year-old BMI and 8-year-old BMI traits are from the same cohort, we incorporate the estimated noise correlation matrix (Figure S4c). Despite significant genetic correlations between the 1-year-old and 8-year-old BMI traits (Figure S4d), FLOW-MR still detected the protective effect of 8-year-old BMI on breast cancer at the p-value threshold 0.001 (Figure 4cd). As the direct effect of 1-year-old BMI on breast cancer is insignificant, the analysis revealed that the protective effect of body size on preventing breast cancer is likely confined to a specific period in childhood.

**Figure 4:**
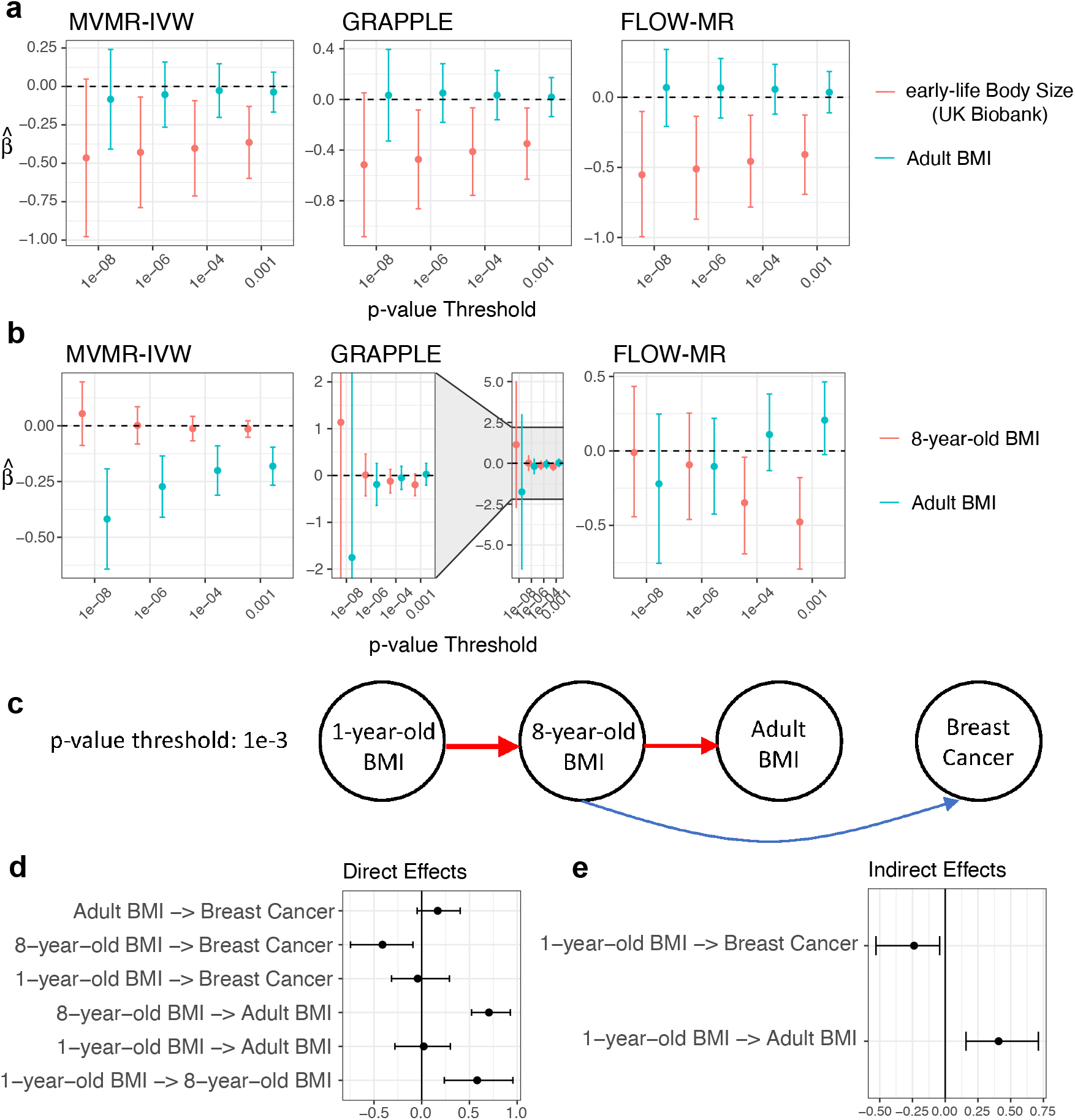
Evaluation of the effect of body size on breast cancer at different ages. a) 95% Confidence / credible intervals of childhood body size (from UK Biobank) and adult BMI on breast cancer risk estimated by MVMR-IVW, GRAPPLE, and FLOW-MR. b) 95% Confidence / credible intervals of childhood BMI and adult BMI on breast cancer risk estimated by these three methods. c) Estimated causal DAG using FLOW-MR at p-value threshold 0.001. The red (blue) arrows indicate significant positive (negative) direct effects at significance level *α* = 0.05. 95% Credible intervals estimated by FLOW-MR are also provided for d) all the direct effects and e) indirect effects of 1-year-old BMI.

Additionally, we found that 1-year-old BMI directly influences 8-year-old BMI but does not have any additional direct effect on adult BMI. Interestingly, the estimated causal effects of 1-year-old BMI on 8-year-old BMI and 8-year-old BMI on adult BMI appear to be of similar magnitude (Figure 4d, Figure S4e). Furthermore, FLOW-MR has the power to detect path-wise indirect causal effects, indicating a significant protective effect of 1-year-old BMI against breast cancer, as well as a positive effect on adult BMI, both of which are mediated through 8-year-old BMI (Figure 4e).

### Temporal causal relationships between high blood pressure, BMI and stroke

High blood pressure is a well-established modifiable risk factor for stroke [30]. Recent univariate MR studies have also suggested that adult BMI may be a potential causal risk factor for stroke and has a significant positive causal effect on high blood pressure in adulthood [14, 15, 31]. Our study aims to disentangle the complex relationships between high blood pressure, BMI, and stroke, focusing on how these causal connections evolve throughout life. We also consider potential confounders, such as lipid traits.

We analyze the causal effects of systolic blood pressure (SBP), BMI, and low-density lipoprotein cholesterol (LDL-C) in both childhood and adulthood on stroke using FLOW-MR. To capture potential causal effects of BMI on SBP, even at the same time point, we introduce additional “time points” (stages) for SBP traits both in childhood and adulthood (Figure 5a). We utilize GWAS summary statistics from the ALSPAC cohort [24] for childhood traits, UK Biobank for adult BMI and SBP, and the Global Lipids Genetics Consortium for adult LDL-C [32]. Stroke data comes from Malik et al. [33], and SNP selection is based on the GERA datasets [34, 35], the GIANT adult BMI dataset, and EGG childhood BMI dataset.

**Figure 5:**
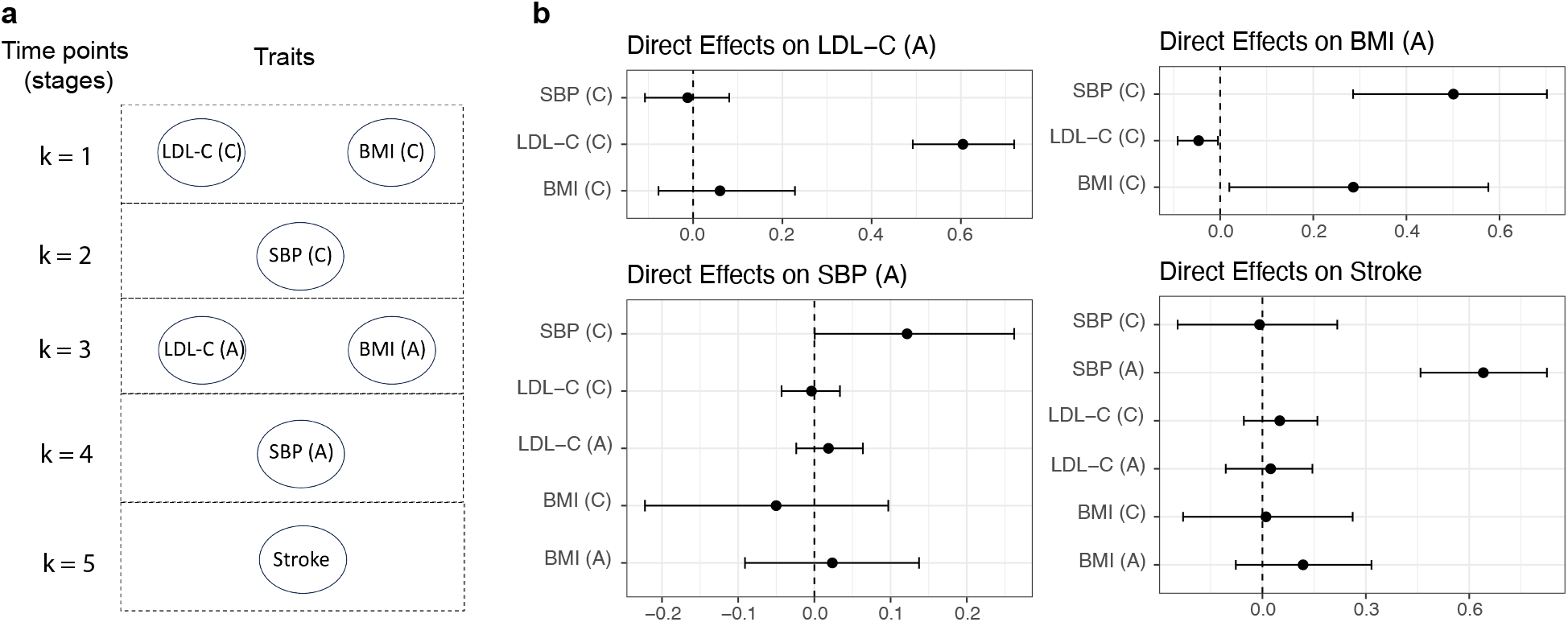
Multivariate mediation analysis on LDL-C, BMI, SBP and stroke. a) Design of the stages. Childhood traits are marked with label (C) and adulthood traits are marked with label (A). b) 95% Credible intervals estimated by FLOW-MR for direct effects on adult traits LDL-C, BMI, SBP and Stroke.

Our analysis reveals a clear positive causal effect of adulthood SBP on stroke, with no significant direct effects from other traits, including childhood traits and adult BMI (Figure 5b, Figure S5a). Interestingly, our findings challenge previous univariate MR results ([14, 15], Figure S5b), showing no evidence of a positive causal effect of adult BMI on adult SBP after accounting for childhood SBP. Further MR analyses using only these three continuous traits also suggest that the previously observed association between adult BMI and SBP may be confounded by childhood SBP (Figure S5c). Additionally, we observe a stronger genetic correlation between childhood SBP and adult BMI than between adult SBP and BMI (Figure S5d). These results imply that the impact of adulthood BMI on SBP in earlier studies may have been overstated due to unaccounted confounding factors.

## Discussion

We propose FLOW-MR using GWAS summary data for life-course Mendelian Randomization, based on a full structural model for all traits. This method allows users to assess time-varying causal effects of heritable risk factors and distinguish between direct and indirect causal effects. By addressing key challenges in life-course MR—the high genetic correlation of the same trait at different ages and the limited cohort size of age-specific GWAS, FLOW-MR demonstrates superior performance in simulation studies. Unlike other methods, FLOW-MR is not prone to biases arising from using weakly associated SNPs as instruments and can efficiently integrate information across all traits.

For the identification of causal effects in life-course MR, a similar argument has been discussed in [7]. They show that to identify the causal direct effects of traits, genetic variants must exert different effects on each exposure in the model, and these effects must be linearly independent, meaning the true SNP-trait association matrix must have full rank. This condition is necessary but not sufficient for identifying all direct and indirect effects. For instance, in univariable MR, pleiotropic effects must adhere to specific assumptions (such as the InSIDE assumption) to ensure the identification of a risk factor’s casual effect. Merely assuming that the pleiotropic effects are not perfectly correlated with the marginal SNP-exposure associations is insufficient for identifying the causal effects. While we allow pervasive pleiotropy in our work, we assume that the direct genetic effects of SNPs are independent across traits, so there is no correlated pleiotropy between the traits. Our sensitivity analysis indicates that, although FLOW-MR models the life-course traits as a whole system, the statistical inference for a particular outcome demonstrates robustness to correlated pleiotropy between other traits (Figure S6). In practice, if confounding pathways are known to exist between traits with correlated pleiotropy, we recommend collecting additional GWAS data for these confounding factors and explicitly adjusting for them using FLOW-MR.

To assess the stability of estimates across different instrument selection criteria, in the paper we employed a range of p-value thresholds for selecting genetic instruments in both simulations and case studies, following the recommendations of [11]. In our simulations, the results remain broadly consistent across the various p-value thresholds. In contrast, larger discrepancies are observed in the real data case studies. This difference may be partly due to the stronger causal signals and the enforced independence of direct genetic effects across traits in the simulation design. Additionally, the number of selected SNPs varies much more substantially in the real data (Table S1), which contributes to differences in statistical power across thresholds.

Finally, as noted by previous studies [6, 36], life-course MR for mediation analysis also faces several potential limitations. One such limitation is the absence of measurements at critical time points when the exposure exerts a substantial causal effect. However, if a missing time point *X*_*k*_ is not a hidden confounder for any pair of subsequent traits 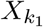 and 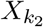, our assumption of independent direct genetic associations remains valid, and the omission of that time point does not bias the estimation of causal effects at observed time points. Another concern is the possibility of reverse causation, where the outcome may causally influence the exposure measured at the latest time point. Although life-course MR typically benefits from the temporal ordering of traits, a challenge arises when the final exposure and outcome are measured concurrently—such as when both are assessed in adulthood. In such cases, if the selected SNPs influence the outcome primarily through their effects on the exposures, MR estimates are generally robust to reverse causality [6, 11, 37]. Our proposed framework inherits this desirable property of MR.

## Methods

### Structural equations on individual-level data

Denote ***Z*** as the vector containing all available genetic variants. Based on the causal DAG (Figure 1), we assume the following additive structural equations for each individual:

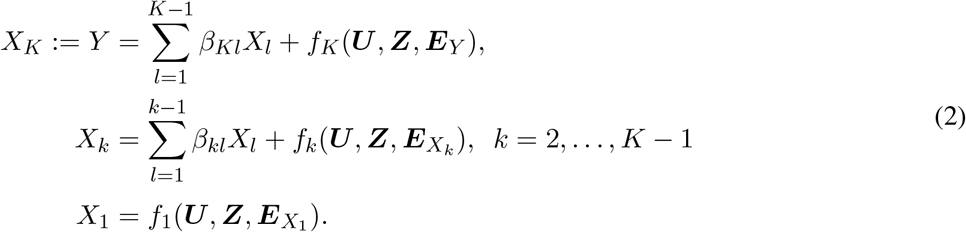

In these equations, the functions *f*_1_(·), …, *f*_*K*_(·) capture the direct genetic and environmental effects on the traits, which may be nonlinear and involve arbitrary interactions. Our primary assumption in (2) is that the causal effect *β*_*kl*_ for any earlier trait *X*_*l*_ (1 ≤ *l* ≤ *K* − 1) on any later trait *X*_*k*_ (*l* + 1 ≤ *k* ≤ *K*) is linear and homogeneous. When the outcome or risk factor traits are binary, we adopt a liability threshold model [16], treating each binary trait *X*_*k*_ as a discretized version of an underlying continuous liability 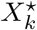, that follows the additive structural equations above (Figure S1). In this context, our method estimates the causal relationships among the latent liabilities. We also further generalize (2) to allow for nonlinear causal effects and interactions among earlier traits, so that our linear model for summary statistics can accommodate more complex causal relationships measured at different time points. Additional details are provided in the SI Text.

Given these structural equations representing the causal relationships among the traits, each *β*_*kl*_ represents the direct causal effect of *X*_*l*_ on *X*_*k*_ whenever *l* < *k*. A path-wise causal effect from *X*_*l*_ to *X*_*k*_ along a path 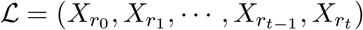 (where 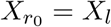 and 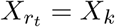 of length *t* is defined as

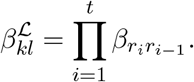

The total effect sums up the effect from *X*_*l*_ to *X*_*k*_ through all directed pathways, and the indirect effect is the difference between the total effect and the direct effects [38, 39]. In other words, the indirect effect 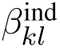 sums all path-wise causal effects of length *t* ≥ 2:

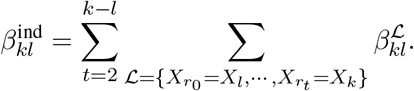

### Model on summary statistics

Denote 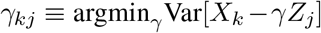 as the marginal association between continous trait *X*_*k*_ and *Z*_*j*_, and let 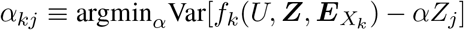 as the projection of the genetic and environmental direct effects of trait *X*_*k*_ onto SNP *Z*_*j*_, which can be heuristically understood as the “direct” genetic effect of SNP *j* on *X*_*k*_. Then by projecting the structural equations (2) onto a single SNP *Z*_*j*_, we can obtain a linear model (1) with measurement error on the GWAS summary statistics (for mathematical details, see SI Text). These equations can be more conveniently expressed in matrix form as:

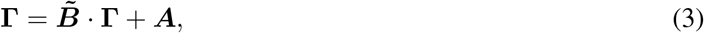

where *P* is the number of SNPs and the matrices are defined as

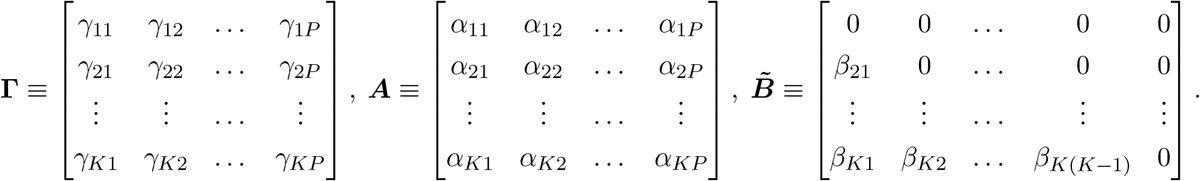

Here, the matrix **Γ** represents the marginal SNP-trait associations, ***A*** denotes the direct effects of the SNPs on each trait and 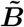 is the matrix of direct causal relationships of earlier traits on later traits that we would like to infer. Equation (3) can be alternatively represented as

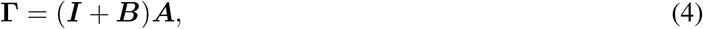

where ***I*** is the *K* × *K* identity matrix, and ***B*** is the lower-triangular matrix that solves 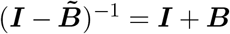. Compared to (3), the parametrization in (4) avoids matrix inversion, making it more amenable for statistical estimation. It is easy to show that ***B*** can be written as a Neumann series: 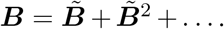 Intuitively, the (*k, l*) entry of the matrix ***B*** denotes the total causal effects of *X*_*l*_ on *X*_*k*_ through all directed pathways [40].

Since the GWAS cohort size is typically large, for each SNP *Z*_*j*_, the summary statistics 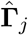 approximately follow a normal distribution 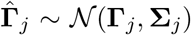, where **Γ**_*j*_ := (*γ*_1*j*_, *γ*_2*j*_, …, *γ*_*Kj*_)^*T*^ is a vector of marginal associations and **Σ**_*j*_ is a covariance matrix obtained from the GWAS standard errors and a correlation matrix that depends on the extent of sample overlap and the correlation between the traits. This correlation matrix is shared across the SNPs and can be estimated from the non-statistically significant GWAS summary statistics using the method described in [11] (SI Text).

### Identification conditions for direct and indirect effects

We provide identification conditions for the matrix 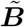 of all direct causal effects from earlier traits to later ones. Once these direct effects are identified, the indirect effects—arising from sums and products of direct effects under our additive structural equations (2)—become identifiable as well.

Under the relationship 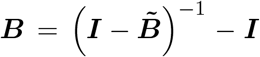, it suffices to identify the lower-triangular matrix ***B***. Assuming an infinitely large cohort, the marginal association matrix **Γ** can be consistently estimated from GWAS data. We then provide conditions on the direct genetic effects ***A*** that make ***B*** uniquely solvable in equation (4).

Specifically, we prove that ***B*** is unique if matrix ***A*** lies in the set

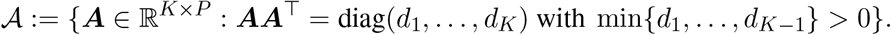

Intuitively, rather than assuming that each trait *k* has its own set of valid SNPs (i.e., there is at least a SNP *j* where *α*_*jk*_ ≠ 0 and *α*_*jk*′_ = 0 for any *k*^′^ ≠ *k*), we require only that the direct genetic effects across traits are mutually orthogonal. Mathematically, this is because (***I*** + ***B***)(***AA***^⊤^)(***I*** + ***B***)^*T*^ provides a unique LDL decomposition of the identifiable matrix **ΓΓ**^⊤^. To make this assumption more realistic, we further can show that when the number of SNPs *P* → ∞, it is sufficient for these direct effects to be merely independent across traits rather than strictly orthogonal—an extension of the well-known InSIDE assumption to multiple traits.

### Selection of genetic instruments

We assume that we can use separate GWAS datasets to obtain p-values for SNP selection, thereby mitigating potential selection biases [11, 41]. Upon selecting the SNPs, another set of GWAS datasets is used to obtain the summary statistics for the exposure and outcome traits. The selection p-value for each SNP is defined as the Bonferroni-corrected p-value, computed from *K* − 1 GWAS summary datasets corresponding to traits *X*_1_ to *X*_*K*−1_. SNPs are subsequently chosen based on the selection p-values using LD clumping [42] (with *r*^2^ cutoff at 0.001), ensuring that selected SNPs are approximately independent. Compared to a stringent p-value threshold (such as 10^−8^) for SNP selection, our method allows for a higher threshold (such as 10^−4^ or 0.01) and uses SNPs that are weakly associated with the exposure traits. This can generally increase the power of MR analysis.

### Model estimation and inference

We use a hierarchical Bayesian framework and Gibbs sampler to infer the direct and indirect causal effects, which can be expressed as functions of the matrix ***B***. In addition to the spike-and-slab prior on the direct genetic effects, we put Gaussian priors on elements of the matrix ***B*** and conjugate priors on the hyperparameters:

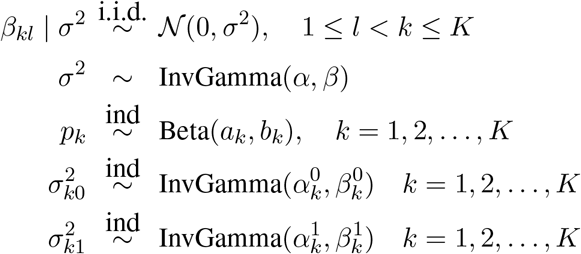

To estimate the above Bayesian hierarchical model, we employ a Gibbs sampler algorithm which can efficiently generate posterior samples when *K* is small. Posterior samples of ***B*** directly give us posterior samples of 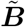, which can then be used to construct credible intervals for any direct and indirect causal effects, or their functions. Additionally, we use a simplified empirical Bayes approach to choose the hyper-parameters 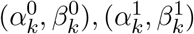 and (*a*_*k*_, *b*_*k*_), recognizing their significant impact on the posterior distributions of ***B***. For further computational and mathematical details, see SI Text.

### Extension to multiple traits at each time point

We also extend our model to accommodate multiple traits at any given time point, incorporating two key assumptions to facilitate the MR analysis. First, we assume that there is no causal effect between traits at the same time point. This assumption avoids the need to identify causal directions between any two traits that coexist temporally. If two sets of traits {*X*_*k*1_, …, *X*_*km*_} and {*X*_*k*(*m*+1)_, …, *X*_*kM*_} at the same time point *k* potentially causally related—with the first set influencing the second—our method can still be applied by introducing an additional “time point” (or stage) *k* + 1 immediately following *k*. The second set of traits is then moved to this added stage, and all subsequent stages are shifted accordingly.

The second assumption is a generalization of the independence assumption on direct genetic effects. Specifically, we require that the direct genetic effects *α*_*kj*_ of any SNP *j* must be independent across all traits, including those measured at the same and at different time points. With these assumptions, we can adapt our model and Gibbs sampler to infer the direct and indirect causal effects.

### Benchmarking methods

We benchmarked the competing methods using publicly available software packages. Specifically, GRAPPLE (https://github.com/jingshuw/GRAPPLE), MVMR-IVW (MVMR; https://github.com/WSpiller/MVMR), MVMR-Egger (MendelianRandomization; https://CRAN.R-project.org/package=MendelianRandomization), MVMR-Robust (https://github.com/aj-grant/robust-mvmr), MVMR-cML (https://github.com/ZhaotongL/MVMR-cML), and MVMR-Horse (https://github.com/aj-grant/mrhorse) were all implemented with their standard procedures recommended in the respective software documentation. We used the same set of selected SNPs for each method.

## Author contributions

J.W. supervised the work. J.W. and Q.Z. formulated the problem. J.W., Z.W. and E.L. developed the algorithm. Z.W. and E.L. implemented the method and performed simulations. Z.W. conducted real data analyses. J.W., Q.Z. and Z.W. wrote the paper.

## Acknowledgments

J.W. is partly supported by the National Science Foundation under grants DMS-2113646 and DMS-2238656. Q.Z. is partly supported by EPSRC grant EP/V049968/1. We acknowledge the University of Chicago’s Research Computing Center for their support of this work.

## Data availability

GWAS summary statistics are all downloaded from public sources. For adult BMI, data is obtained from the GIANT consortium website (https://portals.broadinstitute.org/collaboration/giant/index.php/GIANT_consortium_data_files) and the UK Biobank Neale’s lab (http://www.nealelab.is/uk-biobank/) using phenotype code 21001. Adult lipid traits (LDL-C, HDL-C, and triglycerides) summary statistics are acquired from the GLGC cohort via the GLGC website (https://csg.sph.umich.edu/willer/public/lipids2013/) and from the GERA cohort through GWAS Catalog with study accession numbers GCST007141, GCST007140, and GCST007142. Breast cancer GWAS summary statistics come from GWAS Catalog with study accession number GCST004988, while those for T2D are downloaded from the DIAGRAM website (https://diagram-consortium.org/downloads.html), and for stroke from GWAS Catalog with study accession number GCST006906.

For childhood traits, GWAS summary statistics on 1-year-old BMI and 8-year-old BMI are contributed by the Centre For Diabetes Research, University of Bergen, Norway, and the Norwegian Mother, Father and Child study, downloadable from their website (https://www.fhi.no/en/studies/moba/for-forskere-artikler/gwas-data-from-moba/). Childhood BMI GWAS data from the EGG Consortium can be found at http://egg-consortium.org/childhood-bmi.html. The ALSPAC datasets for childhood lipid traits, SBP, and BMI are obtained from GWAS Catalog with study accession numbers GCST90104679, GCST90104678, GCST90104680, GCST90104683, GCST90104677.

## Code availability

The R package FLOW-MR for conducting our Bayesian mediation MR analysis is publicly available for installation at (https://github.com/ZixuanWu1/FLOW-MR). The code for all experiments, real data analyses, and the reproduction of all figures, along with source data, can be accessed at (https://github.com/ZixuanWu1/FLOW-MR-PAPER).

