## Supplemental Materials for "Causal mediation analysis for time-varying heritable risk factors with Mendelian Randomization"

#### Supplementary Text

#### S1 Additional mathematical details

##### S1.1 Derivation of summary-data linear models

Here, we provide justification of the linear model (1) in the main text from individual-level causal structural equations. We first justify the model under additive structural equations of continuous traits, then extend our justification when any traits are binary. We also interpret model (1) in the main text under general causal relationships among traits.

###### S1.1.1 Derivation under Additive structural equations

Recall that in the Methods section, we assume the following additive structural equations for traits $(X_1, \cdots, X_K)$:

$$\begin{aligned} X_K := Y &= \sum_{l=1}^{K-1} \beta_{Kl} X_l + f_K(\boldsymbol{U}, \boldsymbol{Z}, \boldsymbol{E}_Y), \\ X_k &= \sum_{l=1}^{k-1} \beta_{kl} X_l + f_k(\boldsymbol{U}, \boldsymbol{Z}, \boldsymbol{E}_{X_k}), \quad k = 2, \ldots, K-1 \\ X_1 &= f_1(\boldsymbol{U}, \boldsymbol{Z}, \boldsymbol{E}_{X_1}). \end{aligned} \tag{1}$$

The main assumption is that an earlier trait has a linear and homogenous causal effect on a later trait. We have also defined the marginal associations as

$$\gamma_{kj} \equiv \text{argmin}_{\gamma} \text{Var}[X_k - \gamma Z_j] \tag{2}$$

$$\alpha_{kj} \equiv \text{argmin}_{\alpha} \text{Var}[f_k(\boldsymbol{U}, \boldsymbol{Z}, \boldsymbol{E}_{X_k}) - \alpha Z_j] \tag{3}$$

Then, linearly projecting $X_k$ onto $Z_j$, we have

$$X_k = \gamma_{kj} Z_j + (X_k - \gamma_{kj} Z_j) = \gamma_{kj} Z_j + \epsilon_{kj} \tag{4}$$

where $\text{corr}(Z_j, \epsilon_{kj}) = 0$. On the other hand, we have

$$X_k = \sum_{l=1}^{k-1} \beta_{kl} X_l + \alpha_{kj} Z_j + (f_k(\boldsymbol{U}, \boldsymbol{Z}, \boldsymbol{E}_{\boldsymbol{X}_k}) - \alpha_{kj} Z_j) = \sum_{l=1}^{k-1} \beta_{kl} X_l + \alpha_{kj} Z_j + \tilde{e}_{kj}, \tag{5}$$

where $\text{corr}(Z_j, \tilde{e}_{kj}) = 0$. Substitute (4) into (5) for $l = 1, \ldots, k-1$, we further have

$$X_k = \left(\sum_{l=1}^{k-1} \beta_{kl} \gamma_{lj}\right) Z_j + \alpha_{kj} Z_j + \left(\sum_{l=1}^{k-1} \beta_{kl} \epsilon_{lj}\right) + \tilde{e}_{kj} = \left(\sum_{l=1}^{k-1} \beta_{kl} \gamma_{lj} + \alpha_{kj}\right) Z_j + \left(\sum_{l=1}^{k-1} \beta_{kl} \epsilon_{lj} + \tilde{e}_{kj}\right). \tag{6}$$

Since $\text{corr}(Z_j, (\sum_{l=1}^{k-1} \beta_{kl}\epsilon_{lj} + \tilde{e}_{kj})) = 0$, both (4) and (6) are linear projections of $X_k$ on $Z_j$, thus

$$\gamma_{kj} = \sum_{l=1}^{k-1} \beta_{kl}\gamma_{lj} + \alpha_{kj}, \quad k = 1, \ldots, K \tag{7}$$

##### S1.1.2 Extension to binary traits

When any of the observed traits $X_k$ is binary, we assume a liability threshold model. Specifically, we treat $X_k$ as discretization of a latent continuous trait $X_k^*$, where $X_k^*$ follows the linear structural equations we established before, and $X_k = \mathbf{1}\{X_k^* \geq c_k\}$ for some threshold $c_k \in \mathbb{R}$. For $X_{k'}$ is a continuous trait, then for notation consistency, we define $X_{k'}^* = X_k$. Now let $\gamma_{kj}^*$ denote the marginal association between the latent trait $X_k^*$ and the SNP $Z_j$, as defined in (2). Then following (7) on the continuous traits, we have:

$$\gamma_{kj}^* = \sum_{l=1}^{k-1} \beta_{kl}\gamma_{lj}^* + \alpha_{kj}^*, \quad k = 1, \ldots, K$$

where $\alpha_{kj}^*$ is the marginal association between $f_k(\boldsymbol{U}, \boldsymbol{Z}, \boldsymbol{E}_{\boldsymbol{X}_k^\star})$ and $Z_j$.

Let $\gamma_{kj}$ be the true coefficient regressing $X_k$ on $Z_j$, either using linear or logistic regression, depending on which model the summary statistics for binary traits is used. Then as shown in our recent work [1], under the assumption that each SNP has a negligible effect on $X_k^\star$, $\gamma_{kj}$ and $\gamma_{kj}^\star$ are approximately proportional to each other:

$$\gamma_{kj} \approx s_k \gamma_{kj}^*$$

where $s_k$ is shared across all SNPs and can be calculated given the prevalence $\mathbb{P}(X_k = 1)$ of the binary trait. Thus, the linear model (7) still holds if any of the traits are binary.

##### S1.1.3 Interpretation under general non-linear causal relationship among traits

Following a similar line of reasoning to that presented in Section A.2 of our earlier work [2], we show that the linear relationship (7) approximately holds for a wide range of causal structures among the traits.

Specifically, consider the general causal relationship among traits:

$$\begin{aligned} X_K := Y &= g_K(X_1, \cdots, X_{K-1}, \boldsymbol{U}, \boldsymbol{Z}, \boldsymbol{E}_Y), \\ X_k &= g_k(X_1, \cdots, X_{k-1}, \boldsymbol{U}, \boldsymbol{Z}, \boldsymbol{E}_{X_k}), \quad k = 2, \ldots, K-1 \\ X_1 &= f_1(\boldsymbol{U}, \boldsymbol{Z}, \boldsymbol{E}_{X_1}). \end{aligned} \tag{8}$$

Here, the only assumption we impose is that the functions $g_2, \ldots, g_K$ are differentiable with respect to the trait variables $X_1, \ldots, X_{K-1}$.

We continue to define the marginal associations $\gamma_{kj}$ as (2) so that each $X_k = \gamma_{kj}Z_j + \epsilon_{kj}$. Again, we assume that a single SNP has negligible association on a complex trait, so that $\gamma_{kj} = o(1)$ for any $k$ and $j$, then for any $k > 1$ and $j$,

$$\begin{aligned} g_k(X_1, \cdots, X_{k-1}, \boldsymbol{U}, \boldsymbol{Z}, \boldsymbol{E}_{X_k}) &= g_k(\gamma_{1j}Z_j + \epsilon_{1j}, \cdots, \gamma_{(k-1)j}Z_j + \epsilon_{(k-1)j}, \boldsymbol{U}, \boldsymbol{Z}, \boldsymbol{E}_{X_k}) \\ \approx g_k(\epsilon_{1j}, \cdots, \epsilon_{(k-1)j}, \boldsymbol{U}, \boldsymbol{Z}, \boldsymbol{E}_{X_k}) &+ \sum_{l=1}^{k-1} \frac{\partial g_k}{\partial x_l}(X_1, \cdots, X_{k-1}, \boldsymbol{U}, \boldsymbol{Z}, \boldsymbol{E}_{X_k})\gamma_{lj}Z_j. \end{aligned}$$

Now we define the causal effect of trait $X_l$ on a later trait $X_k$ by the average derivative effect (ADE) [3], defined as

$$\beta_{kl} = \mathbb{E}\left[\frac{\partial g_k}{\partial x_l}(X_1, \cdots, X_{k-1}, \boldsymbol{U}, \boldsymbol{Z}, \boldsymbol{E}_{X_k})\right].$$

Then because $\frac{\partial g_k}{\partial x_l}(X_1, \cdots, X_{k-1}, \boldsymbol{U}, \boldsymbol{Z}, \boldsymbol{E}_{X_k})$ is still a complex trait, we should have

$$\text{Cov}\left[\frac{\partial g_k}{\partial x_l}(X_1, \cdots, X_{k-1}, \boldsymbol{U}, \boldsymbol{Z}, \boldsymbol{E}_{X_k}), Z_j\right] = o(1)$$

for most SNPs. As a consequence, we have

$$\text{Cov}\left[\left(\frac{\partial g_k}{\partial x_l}(X_1,\cdots,X_{k-1},\boldsymbol{U},\boldsymbol{Z},\boldsymbol{E}_{X_k})-\beta_{kl}\right)Z_j,Z_j\right]=o(1).$$

Thus we have for any $l<k$,

$$\frac{\partial g_k}{\partial x_l}(X_1,\cdots,X_{k-1},\boldsymbol{U},\boldsymbol{Z},\boldsymbol{E}_{X_k})\gamma_{lj}Z_j=\beta_{kl}\gamma_{lj}Z_j+o(1)\gamma_{lj}Z_j+\tilde{e}_j$$

where $\text{cor}(\tilde{e}_j,Z_j)=0$.

Further, we can define

$$\alpha_{kj}\equiv\text{argmin}_{\alpha}\text{Var}\left[g_k(\epsilon_{1j},\cdots,\epsilon_{(k-1)j},\boldsymbol{U},\boldsymbol{Z},\boldsymbol{E}_{X_k})-\alpha Z_j\right],$$

thus we still have

$$\gamma_{kj}\approx\sum_{l=1}^{k-1}\beta_{kl}\gamma_{lj}+\alpha_{kj},\quad k=1,\ldots,K$$

for any $j$.

#### S1.2 Noise correlation estimation

As shown in [4], for any risk factor $k$ and $l$ we have

$$\text{Corr}(\hat{\gamma}_{kj},\hat{\gamma}_{lj})\approx\frac{N_{kl}}{\sqrt{N_kN_l}}\text{Corr}(X_{ks},X_{ls}),\tag{9}$$

where $N_k$ and $N_l$ are the sample sizes of the risk factor $X_k$ and $X_l$, $N_{kl}$ is number of overlapping individuals, and $X_{ks}$ denotes the measure of $X_k$ for individuals $s$. The correlation of $X_k$ and $X_l$ of any shared sample is $\text{Corr}[X_{ks},X_{ls}]$. Thus, the correlation between $\hat{\gamma}_{kj}$ is approximately the same for all SNPs. As a consequence, we assume

$$\begin{pmatrix}\hat{\gamma}_{1j}\\ \hat{\gamma}_{2j}\\ \vdots\\ \hat{\gamma}_{Kj}\end{pmatrix}\sim\mathcal{N}\left(\begin{pmatrix}\gamma_{1j}\\ \gamma_{2j}\\ \vdots\\ \gamma_{Kj}\end{pmatrix},\begin{pmatrix}\delta_{1j}&&&\\&\delta_{2j}&&\\&&\ddots&\\&&&\delta_{Kj}\end{pmatrix}\boldsymbol{R}\begin{pmatrix}\delta_{1j}&&&\\&\delta_{2j}&&\\&&\ddots&\\&&&\delta_{Kj}\end{pmatrix}\right).$$

Here $\delta_{kj}^2$ is the variance on the estimator $\hat{\gamma}_{kj}$ given as GWAS summary statistics and $\boldsymbol{R}$ is the unknown shared correlation matrix of the SNP associations.

To estimate $\boldsymbol{R}$, we choose SNPs where $\gamma_{kj}=0$ for all $k$ so that we can estimate the shared correlations using the sample correlation of the chosen SNPs. To do this we select SNPs with p-values $p_j\geq 0.5$ in all selection files.

#### S1.3 Formal definition and proof for identifiability

In this section, we provide a formal mathematical statement for identifying the causal effect matrix $\tilde{\boldsymbol{B}}$ in the linear model:

$$\boldsymbol{\Gamma}_P=\tilde{\boldsymbol{B}}\cdot\boldsymbol{\Gamma}_P+\boldsymbol{A}_P\tag{10}$$

where

$$\boldsymbol{\Gamma}_P\equiv\begin{bmatrix}\gamma_{11}&\gamma_{12}&\cdots&\gamma_{1P}\\ \gamma_{21}&\gamma_{22}&\cdots&\gamma_{2P}\\ \vdots&\vdots&\cdots&\vdots\\ \gamma_{K1}&\gamma_{K2}&\cdots&\gamma_{KP}\end{bmatrix},\ \boldsymbol{A}_P\equiv\begin{bmatrix}\alpha_{11}&\alpha_{12}&\cdots&\alpha_{1P}\\ \alpha_{21}&\alpha_{22}&\cdots&\alpha_{2P}\\ \vdots&\vdots&\cdots&\vdots\\ \alpha_{K1}&\alpha_{K2}&\cdots&\alpha_{KP}\end{bmatrix},\ \tilde{\boldsymbol{B}}_P\equiv\begin{bmatrix}0&0&\cdots&0&0\\ \beta_{21}&0&\cdots&0&0\\ \vdots&\vdots&\cdots&\vdots&\vdots\\ \beta_{K1}&\beta_{K2}&\cdots&\beta_{K(K-1)}&0\end{bmatrix}.$$

For identifiability, we assume that the cohort size is unlimited so that the matrix of marginal associations $\boldsymbol{\Gamma}_P$ is always identifiable.

In the main text, it was highlighted that the causal effect matrix $\tilde{\boldsymbol{B}}$ is not identifiable if we only have genetic instruments for the earliest exposure trait $X_1$. Here we give a simple example to illustrate how the direct and indirect effects can be not inseparable in this case. Suppose

$$X_1 = \sum_{j=1}^{P} \alpha_j Z_j,$$

$$X_2 = \beta_1 X_1 = \sum_{j=1}^{P} \beta_1 \alpha_j Z_j,$$

$$Y = \beta_2 X_1 + \beta_3 X_2 = \sum_{j=1}^{P} (\beta_2 + \beta_1\beta_3)\alpha_j Z_j,$$

where $\boldsymbol{Z} = (Z_1, Z_2, \ldots, Z_P)$ are independent SNPs. In this setup, all the SNPs are valid instruments of the earlier trait $X_1$. Suppose we have infinite sample size, thus the marginal associations $(\alpha_1, \alpha_2, \ldots, \alpha_P)$, $(\beta_1\alpha_1, \beta_1\alpha_2, \cdots, \beta_1\alpha_P)$ and $((\beta_2 + \beta_1\beta_3)\alpha_1, (\beta_2 + \beta_1\beta_3)\alpha_2, \cdots, (\beta_2 + \beta_1\beta_3)\alpha_P)$ are all known. Then, $\beta_1$ and $\beta_2 + \beta_1\beta_3$ can be identified as they are ratios of these marginal associations. However, for any $\beta_3' \in \mathbb{R}$, one can define $\beta_2' = \beta_2 - \beta_1(\beta_3' - \beta_3)$, so that $(\beta_2' + \beta_1\beta_3') = (\beta_2 + \beta_1\beta_3)$. Hence in this case the direct effects and indirect effects of $X_1$ on $Y$ are not identifiable.

Next we provide a sufficient condition for the identifiability of the causal effect matrix $\tilde{\boldsymbol{B}}$. Recall that we can define $\boldsymbol{B} = \left(\boldsymbol{I} - \tilde{\boldsymbol{B}}\right)^{-1} - \boldsymbol{I}$ which is also a lower triangular matrix, then we obtain

$$\boldsymbol{\Gamma}_P = (\boldsymbol{I} + \boldsymbol{B})\boldsymbol{A}_P \tag{11}$$

where $\boldsymbol{\Gamma}_P$ is the matrix of true SNP-trait marginal associations that is identifiable from the GWAS summary statistics. There is a one-to-one correspondence between $\boldsymbol{B}$ and $\tilde{\boldsymbol{B}}$ as $\tilde{\boldsymbol{B}} = \boldsymbol{I} - (\boldsymbol{I} + \boldsymbol{B})^{-1}$, thus we only need to prove the identifiability of $\boldsymbol{B}$ given (11).

To begin with, we define the following sets:

$$\mathcal{A}_P := \{\boldsymbol{A}_P \in \mathbb{R}^{K \times P} : \boldsymbol{A}_P \boldsymbol{A}_P^\top = \mathrm{diag}(d_1, \ldots, d_K) \text{ with } \min\{d_1, \ldots, d_{K-1}\} > 0\}$$

Intuitively, the condition $\boldsymbol{A}_P \in \mathcal{A}_P$ implies that the direct genetic effects of the $P$ SNPs are orthogonal across different traits, and for all exposure traits $X_k$ with $k <= K - 1$, there needs to be at least one $j$ where $\alpha_{kj} \neq 0$. We now state the following identifiability result, showing that $A_p \in \mathcal{A}_P$ is a sufficient condition for identifying $\boldsymbol{B}$:

**Theorem S1.1.** *Let $\boldsymbol{B}$ and $\boldsymbol{B}'$ be lower-triangular matrices with zero diagonals. For any $\boldsymbol{A}_P, \boldsymbol{A}_P' \in \mathcal{A}_P$, we have $\boldsymbol{\Gamma}_P = (\boldsymbol{I} + \boldsymbol{B})\boldsymbol{A}_P = (\boldsymbol{I} + \boldsymbol{B}')\boldsymbol{A}_P'$ if and only if $\boldsymbol{B} = \boldsymbol{B}'$ and $\boldsymbol{A}_P = \boldsymbol{A}_P'$.*

*Proof.* Let $\boldsymbol{D}_P = \boldsymbol{A}_P \boldsymbol{A}_P^\top = \mathrm{diag}(d_1, d_2, \cdots, d_K)$ and $\boldsymbol{D}_P' = \boldsymbol{A}_P'(\boldsymbol{A}_P')^\top = \mathrm{diag}(d_1', d_2', \cdots, d_K')$.

Assume $(\boldsymbol{I} + \boldsymbol{B})\boldsymbol{A}_P = (\boldsymbol{I} + \boldsymbol{B}')\boldsymbol{A}_P'$. Note

$$((\boldsymbol{I} + \boldsymbol{B})\boldsymbol{A}_P)\left((\boldsymbol{I} + \boldsymbol{B})\boldsymbol{A}_P\right)^\top = (\boldsymbol{I} + \boldsymbol{B})\boldsymbol{D}_P(\boldsymbol{I} + \boldsymbol{B})^\top$$

$$\left((\boldsymbol{I} + \boldsymbol{B}')\boldsymbol{A}_P'\right)\left((\boldsymbol{I} + \boldsymbol{B}')\boldsymbol{A}_P'\right)^\top = (\boldsymbol{I} + \boldsymbol{B}')\boldsymbol{D}_P'(\boldsymbol{I} + \boldsymbol{B}')^\top$$

Therefore

$$\boldsymbol{M} \triangleq \boldsymbol{\Gamma}\boldsymbol{\Gamma}^\top = (\boldsymbol{I} + \boldsymbol{B})\boldsymbol{D}_P(\boldsymbol{I} + \boldsymbol{B})^\top = (\boldsymbol{I} + \boldsymbol{B}')\boldsymbol{D}_P'(\boldsymbol{I} + \boldsymbol{B}')^\top.$$

Since both $\boldsymbol{I} + \boldsymbol{B}$ and $\boldsymbol{I} + \boldsymbol{B}'$ are lower unitriangular matrices, they become two LDL decompositions of the identifiable matrix $\boldsymbol{M}$. We basically need to show that the LDL decomposition of $\boldsymbol{M}$ is unique.

In particular we show that if for $i \geq j$, we have

$$\left[(\boldsymbol{I} + \boldsymbol{B})\boldsymbol{D}_P(\boldsymbol{I} + \boldsymbol{B})^\top\right]_{ij} = \sum_{k=1}^{j} (\boldsymbol{I} + \boldsymbol{B})_{ik} d_k (\boldsymbol{I} + \boldsymbol{B})_{jk} = \sum_{k=1}^{j} (\boldsymbol{I} + \boldsymbol{B}')_{ik} d_k' (\boldsymbol{I} + \boldsymbol{B}')_{jk},$$

then $\beta_{ij} = \beta_{ij}'$ and $d_j = d_j'$ for $j \leq K - 1$. We prove by induction.

**Base Case:** For $j = 1$,

$$\left[(\boldsymbol{I}+\boldsymbol{B})\boldsymbol{D}_P(\boldsymbol{I}+\boldsymbol{B})^\top\right]_{i1} = (\boldsymbol{I}+\boldsymbol{B})_{i1}d_1.$$

Similarly,

$$\left[(\boldsymbol{I}+\boldsymbol{B}')\boldsymbol{D}'_P(\boldsymbol{I}+\boldsymbol{B}')^\top\right]_{i1} = (\boldsymbol{I}+\boldsymbol{B}')_{i1}d'_1.$$

Since $(\boldsymbol{I}+\boldsymbol{B})_{11} = (\boldsymbol{I}+\boldsymbol{B}')_{11} = 1$, we conclude $d_1 = d'_1$. Since $d_1 > 0$, it follows that $\beta_{i1} = \beta'_{i1}$ for $i \geq 1$.

**Inductive Step:** Assume $\beta_{ij} = \beta'_{ij}$ and $d_j = d'_j$ for $j \leq m-1$ (where $m \leq K-1$). For $j = m$, with $i \geq m$, we have

$$\begin{aligned}
\left[(\boldsymbol{I}+\boldsymbol{B})\boldsymbol{D}_P(\boldsymbol{I}+\boldsymbol{B})^\top\right]_{im} &= \sum_{k=1}^{m}(\boldsymbol{I}+\boldsymbol{B})_{ik}d_k(\boldsymbol{I}+\boldsymbol{B})_{mk} \\
&= \sum_{k=1}^{m-1}(\boldsymbol{I}+\boldsymbol{B})_{ik}d_k(\boldsymbol{I}+\boldsymbol{B})_{mk} + (\boldsymbol{I}+\boldsymbol{B})_{im}d_m, \\
\left[(\boldsymbol{I}+\boldsymbol{B}')\boldsymbol{D}'_P(\boldsymbol{I}+\boldsymbol{B}')^\top\right]_{im} &= \sum_{k=1}^{m}(\boldsymbol{I}+\boldsymbol{B}')_{ik}d'_k(\boldsymbol{I}+\boldsymbol{B}')_{mk} \\
&= \sum_{k=1}^{m-1}(\boldsymbol{I}+\boldsymbol{B}')_{ik}d'_k(\boldsymbol{I}+\boldsymbol{B}')_{mk} + (\boldsymbol{I}+\boldsymbol{B}')_{im}d'_m.
\end{aligned}$$

By the induction hypothesis, for $i \geq m$,

$$(\boldsymbol{I}+\boldsymbol{B})_{im}d_m = (\boldsymbol{I}+\boldsymbol{B}')_{im}d'_m.$$

Since $(\boldsymbol{I}+\boldsymbol{B})_{mm} = (\boldsymbol{I}+\boldsymbol{B}')_{mm} = 1$, it follows that $d_m = d'_m$, and since $d_m > 0$, we conclude $\beta_{im} = \beta'_{im}$ for $i \geq m$. This completes the induction.

Finally, we have

$$\boldsymbol{A}_P = (\boldsymbol{I}+\boldsymbol{B})^{-1}\mathbb{E}[\hat{\boldsymbol{\Gamma}}_P] = (\boldsymbol{I}+\boldsymbol{B}')^{-1}\boldsymbol{\Gamma}_P = \boldsymbol{A}'_P.$$

□

We also extend the scope of Theorem S1.1 by relaxing the orthogonality condition on the direct genetic effects. Instead of requiring these effects to be exactly orthogonal, we now only assume that as the number of SNPs $P \to \infty$, the direct genetic effects behave as if they are "independent" across traits. This generalization broadens the applicability of the theoretical guarantees presented in Theorem S1.1.

To begin with, define $\boldsymbol{D}_P = \boldsymbol{A}_P\boldsymbol{A}_P^\top \in \mathbb{R}^{K\times K}$. We say a sequence of matrices $(\boldsymbol{A}_P \in \mathbb{R}^{K\times P})_{P=1}^\infty$ is asymptotically orthogonal if there exists sequence $(a_P, b_P, c_P)_{P=1}^\infty$ such that

$$\min_{1\leq i=j<K}\{(\boldsymbol{D}_P)_{i,j}\} \geq a_P, \quad \max_{1\leq i=j\leq K}\{(\boldsymbol{D}_P)_{i,j}\} \leq b_P, \quad \max_{i\neq j}\{(\boldsymbol{D}_P)_{i,j}\} \leq c_P,$$

where $(a_P, b_P, c_P)_{P=1}^\infty$ satisfy

$$\lim_{P\to\infty}\frac{c_P}{a_P} = 0, \quad \liminf_{P\to\infty}\frac{a_P}{b_P} = c > 0.$$

Intuitively, we require the direct genetic effects to have vanishing correlations as $P \to \infty$. Then we formally state the identifiability result in the asymptotic regime.

**Corollary S1.1.1.** *Let $\boldsymbol{B}$ and $\boldsymbol{B}'$ be lower-triangular matrices with zero diagonals. For any asymptotic orthogonal sequences $(\boldsymbol{A}_P)_{P=1}^\infty, (\boldsymbol{A}'_P)_{P=1}^\infty$ associated with the sequence $(a_P, b_P, c_P)_{P=1}^\infty$, we have $\boldsymbol{\Gamma}_P = (\boldsymbol{I}+\boldsymbol{B})\boldsymbol{A}_P = (\boldsymbol{I}+\boldsymbol{B}')\boldsymbol{A}'_P$ for all $P$ if and only if $\boldsymbol{B} = \boldsymbol{B}'$.*

*Proof.* Note

$$\liminf_{P\to\infty} \frac{1}{a_P}\boldsymbol{\Gamma}_P\boldsymbol{\Gamma}_P^\top = \liminf_{P\to\infty} \frac{1}{a_P}(\boldsymbol{I}+\boldsymbol{B})\boldsymbol{D}_P(\boldsymbol{I}+\boldsymbol{B})^\top = (\boldsymbol{I}+\boldsymbol{B})\left(\liminf_{P\to\infty} \frac{1}{a_P}\boldsymbol{D}_P\right)(\boldsymbol{I}+\boldsymbol{B})^\top$$

Similarly

$$\liminf_{P\to\infty} \frac{1}{a_P}\boldsymbol{\Gamma}_P\boldsymbol{\Gamma}_P^\top = (\boldsymbol{I}+\boldsymbol{B}')\left(\liminf_{P\to\infty} \frac{1}{a_P}\boldsymbol{D}'_P\right)(\boldsymbol{I}+\boldsymbol{B}')^\top$$

By assumption, $\boldsymbol{D}^\star := \left(\liminf_{P\to\infty} \frac{1}{a_P}\boldsymbol{D}_P\right)$ and $(\boldsymbol{D}')^\star := \left(\liminf_{P\to\infty} \frac{1}{a_P}\boldsymbol{D}'_P\right)$ are diagonal with $\min\{\boldsymbol{D}^\star_{ii}, i=1,\cdots,K-1\} > 0$ and $\min\{(\boldsymbol{D}')^\star_{ii}, i=1,\cdots,K-1\} > 0$. Following the proof of Theorem S1.1, we know $\liminf_{P\to\infty} \frac{1}{a_P}\boldsymbol{\Gamma}_P\boldsymbol{\Gamma}_P^\top$ admits a unique LDL decomposition, which indicates that $\boldsymbol{B} = \boldsymbol{B}'$. □

#### S1.4 Details of the Gibbs Sampler

We observe GWAS summary statistics, $\hat{\gamma}_{kj}$, of marginal associations between SNPs and traits, with standard errors $\delta_{kj}$. Denote $\boldsymbol{\Delta}_j = \text{diag}(\delta_{1j}, \ldots \delta_{Kj})$. Denote the correlation matrix betwen trait-association for all SNPs by $\boldsymbol{R}$.

The model is

$$\hat{\boldsymbol{\Gamma}} = (\boldsymbol{I}+\boldsymbol{B})\boldsymbol{A} + \boldsymbol{\epsilon}$$

$$\begin{aligned}
\boldsymbol{\epsilon_{*,j}} &\sim \mathcal{N}(\boldsymbol{0}, \boldsymbol{\Sigma}_j := \boldsymbol{\Delta}_j\boldsymbol{R}\boldsymbol{\Delta_j}), \text{ with } \boldsymbol{\epsilon_{*,j}} \perp\!\!\!\perp \{\boldsymbol{\epsilon_{*,j'}} : j' \neq j\} \\
\beta_{kl} &\overset{\text{iid}}{\sim} \mathcal{N}(0,\sigma^2) \text{ for } k > l, 0 \text{ otherwise} \\
\sigma^2 &\overset{\text{iid}}{\sim} \text{InvGamma}(\alpha^B, \beta^B) \\
\alpha_{kj} &\overset{\text{iid}}{\sim} \begin{cases} \mathcal{N}(0,\sigma_{k0}^2), & \text{if } z_{kj} = 0 \\ \mathcal{N}(0,\sigma_{k1}^2), & \text{if } z_{kj} = 1 \end{cases} \\
\sigma_{k0} &\overset{\text{iid}}{\sim} \text{InvGamma}(\alpha_k^0, \beta_k^0) \\
\sigma_{k1} &\overset{\text{iid}}{\sim} \text{InvGamma}(\alpha_k^1, \beta_k^1) \\
z_{kj} &\overset{\text{iid}}{\sim} \text{Bernoulli}(p_k) \\
p_k &\overset{\text{iid}}{\sim} \text{Beta}(a_k, b_k)
\end{aligned}$$

We have $K$ traits and $P$ genes. The prior parameters are $\alpha^B$, $\beta^B$, $\alpha_k^0$, $\beta_k^0$, $\alpha_k^1$, $\beta_k^1$, $a_k$ and $b_k$. Also let

$$\begin{aligned}
\boldsymbol{Z} &= (z_{kj})_{K\times P} \\
\boldsymbol{p} &= (p_1, p_2, \ldots, p_K)^T \\
\boldsymbol{\sigma_0} &= (\sigma_{10}, \sigma_{20}, \ldots, \sigma_{K0})^T \\
\boldsymbol{\sigma_1} &= (\sigma_{11}, \sigma_{21}, \ldots, \sigma_{K1})^T
\end{aligned}$$

##### S1.4.1 Notation

For a $K \times P$ matrix $\boldsymbol{M}$, let

- $\boldsymbol{M}_{i,<k}$ denotes a row vector of all elements of $\boldsymbol{M}$ in the $i^{th}$ row and in columns less than $k$
- $\boldsymbol{M}_{i,*}$ denotes a whole row, and $\boldsymbol{M}_{*,j}$ a column.

- $\text{vec}(\boldsymbol{M})$ denotes the columnwise vectorization of $\boldsymbol{M}$, i.e.,

$$\text{vec}(\boldsymbol{M}) = \begin{pmatrix} \boldsymbol{M}_{*,1} \\ \boldsymbol{M}_{*,2} \\ \vdots \\ \boldsymbol{M}_{*,P} \end{pmatrix}.$$

- For $K = P$, define $\text{LTvec}(\boldsymbol{M})$ to be the *lower-triangular vectorization* of $\boldsymbol{M}$ (excluding the diagonal):

$$\text{LTvec}(\boldsymbol{M}) := \begin{pmatrix} M_{2,1} \\ (\boldsymbol{M}_{3,<3})^T \\ \vdots \\ (\boldsymbol{M}_{K-1,<K-1})^T \\ (\boldsymbol{M}_{K,<K})_T \end{pmatrix},$$

  which is a $\binom{K}{2}$-dimensional vector.

For $\boldsymbol{v} \in \mathbb{R}^K$, define

$$\text{LTstack}(\boldsymbol{v}) := \begin{pmatrix} 0 & v_1 & 0 & 0 & \cdots & 0 \\ 0 & 0 & v_1 & 0 & & \\ 0 & 0 & v_2 & 0 & & \\ 0 & 0 & 0 & v_1 & & \\ 0 & 0 & 0 & v_2 & & \\ 0 & 0 & 0 & v_3 & & \\ \vdots & \vdots & \vdots & \vdots & & \vdots \\ 0 & 0 & 0 & 0 & 0 & v_1 \\ \vdots & \vdots & \vdots & \vdots & & \vdots \\ 0 & 0 & 0 & 0 & & v_{k-2} \\ 0 & 0 & 0 & 0 & \cdots & v_{k-1} \end{pmatrix} = \begin{pmatrix} 0 & \boldsymbol{v}_{<2} & 0 & & \cdots & 0 \\ \mathbf{0}_2 & \mathbf{0}_2 & \boldsymbol{v}_{<3} & & \cdots & \vdots \\ \vdots & \vdots & & \ddots & & \\ \mathbf{0}_{K-2} & \mathbf{0}_{K-2} & & & \boldsymbol{v}_{<K-1} & \mathbf{0}_{K-2} \\ \mathbf{0}_{K-1} & \mathbf{0}_{K-1} & \cdots & & \mathbf{0}_{K-1} & \boldsymbol{v}_{<K} \end{pmatrix},$$

which has dimensions $\binom{K}{2} \times K$. Here $\mathbf{0}_k$ denotes the zero vector of dimension $k$.

Hence, if $\boldsymbol{M}$ is square, lower-triangular with zero diagonal,

$$\boldsymbol{M}\boldsymbol{v} = \left[\text{LTvec}(\boldsymbol{M})^T \text{LTstack}(\boldsymbol{v})\right]^T.$$

Finally, for random variables $U$ and $V$, let $p(U|V)$ denote the conditional density of $U$ given $V$.

##### S1.4.2 Updating $\boldsymbol{B}$

With prior:

$$\text{LTvec}(\boldsymbol{B}) \sim N(\mathbf{0}, \sigma^2 \boldsymbol{I}), \tag{12}$$

likelihood (noise distribution):

$$\boldsymbol{\epsilon}_{*,j} \overset{\text{iid}}{\sim} N(\mathbf{0}, \boldsymbol{\Sigma}_j),$$

so that

$$\text{vec}(\boldsymbol{\epsilon}) \sim \mathcal{N}\left(\mathbf{0}, \boldsymbol{\Sigma} := \begin{pmatrix} \boldsymbol{\Sigma}_1 & \mathbf{0} & \cdots & \mathbf{0} \\ \mathbf{0} & \boldsymbol{\Sigma}_2 & \cdots & \mathbf{0} \\ \vdots & & \ddots & \vdots \\ \mathbf{0} & \mathbf{0} & \cdots & \boldsymbol{\Sigma}_P \end{pmatrix}\right) \tag{13}$$

We have that

$$\begin{aligned}
&\hat{\boldsymbol{\Gamma}} = \boldsymbol{A} + \boldsymbol{B}\boldsymbol{A} + \boldsymbol{\epsilon} \\
\implies &\hat{\boldsymbol{\Gamma}} - \boldsymbol{A} = \boldsymbol{B}\boldsymbol{A} + \boldsymbol{\epsilon} \\
\implies &\text{vec}(\hat{\boldsymbol{\Gamma}} - \boldsymbol{A})^T = \text{LTvec}(\boldsymbol{B})^T \tilde{\boldsymbol{A}} + \text{vec}(\boldsymbol{\epsilon})^T \\
\implies &\text{vec}(\hat{\boldsymbol{\Gamma}} - \boldsymbol{A}) = \tilde{\boldsymbol{A}}^T \text{LTvec}(\boldsymbol{B}) + \text{vec}(\boldsymbol{\epsilon})
\end{aligned}$$

Where

$$\tilde{\boldsymbol{A}} := \begin{pmatrix} \text{LTstack}(\boldsymbol{A}_{*,1}) & \text{LTstack}(\boldsymbol{A}_{*,2}) & \cdots & \text{LTstack}(\boldsymbol{A}_{*,P}) \end{pmatrix}$$

From Theorem 2.2 in [5], we know the posterior is

$$\begin{aligned}
\text{LTvec}(\boldsymbol{B}) &\sim N(\boldsymbol{m}, \boldsymbol{C}), \\
\boldsymbol{C} &= (\tilde{\boldsymbol{A}}\boldsymbol{\Sigma}^{-1}\tilde{\boldsymbol{A}}^T + \sigma^{-2}\boldsymbol{I})^{-1}, \\
\boldsymbol{m} &= \boldsymbol{C}\tilde{\boldsymbol{A}}\boldsymbol{\Sigma}^{-1}\,\text{vec}(\hat{\boldsymbol{\Gamma}} - \tilde{\boldsymbol{A}}).
\end{aligned}$$

###### S1.4.3 Updating $\sigma^2, \boldsymbol{\sigma}_0, \boldsymbol{\sigma}_1$

First observe that conditioning on $\boldsymbol{B}$, we have

$$\sigma \perp\!\!\!\perp (\boldsymbol{A}, \boldsymbol{Z}, \boldsymbol{p}, \boldsymbol{\sigma_0}, \boldsymbol{\sigma_1}, \hat{\boldsymbol{\Gamma}}) \mid \boldsymbol{B}$$

(as $\sigma$ is independent of $\boldsymbol{A}, \boldsymbol{Z}, \boldsymbol{p}, \boldsymbol{\sigma_0}, \boldsymbol{\sigma_1}$ and depends on $\hat{\boldsymbol{\Gamma}}$ only through $\boldsymbol{B}$)
If follows that

$$\begin{aligned}
p(\sigma|\boldsymbol{B}, \boldsymbol{A}, \boldsymbol{Z}, \boldsymbol{p}, \boldsymbol{\sigma_0}, \boldsymbol{\sigma_1}, \hat{\boldsymbol{\Gamma}}) &= \frac{p(\sigma, \boldsymbol{A}, \boldsymbol{Z}, \boldsymbol{p}, \boldsymbol{\sigma_0}, \boldsymbol{\sigma_1}, \hat{\boldsymbol{\Gamma}}|\boldsymbol{B})}{p(\boldsymbol{A}, \boldsymbol{Z}, \boldsymbol{p}, \boldsymbol{\sigma_0}, \boldsymbol{\sigma_1}, \hat{\boldsymbol{\Gamma}}|\boldsymbol{B})} \\
&= \frac{p(\boldsymbol{A}, \boldsymbol{Z}, \boldsymbol{p}, \boldsymbol{\sigma_0}, \boldsymbol{\sigma_1}, \hat{\boldsymbol{\Gamma}}|\boldsymbol{B})p(\sigma|\boldsymbol{B})}{p(\boldsymbol{A}, \boldsymbol{Z}, \boldsymbol{p}, \boldsymbol{\sigma_0}, \boldsymbol{\sigma_1}, \hat{\boldsymbol{\Gamma}}|\boldsymbol{B})} \\
&= p(\sigma|\boldsymbol{B})
\end{aligned}$$

So if suffices to consider $p(\sigma|\boldsymbol{B})$.

We have prior:

$$\sigma^2 \sim \text{InvGamma}(\alpha^B, \beta^B),$$

and likelihood for the lower-triangular portion:

$$\beta_{kl} \overset{\text{iid}}{\sim} \mathcal{N}(0, \sigma^2),$$

so we have posterior:

$$\sigma^2 \sim \text{InvGamma}\left(\alpha^B + \frac{n}{2},\ \beta^B + \frac{1}{2}\|\boldsymbol{B}\|_F^2\right),$$

where $n = K(K-1)/2$ is the number of degrees of freedom in $\boldsymbol{B}$ and $\|\cdot\|_F$ is the Frobenius norm.

Similarly for $\boldsymbol{\sigma_0}$ and $\boldsymbol{\sigma_1}$, we have

$$\boldsymbol{\sigma_0} \perp\!\!\!\perp (\boldsymbol{B}, \boldsymbol{p}, \sigma, \boldsymbol{\sigma_1}, \hat{\boldsymbol{\Gamma}}) \mid (\boldsymbol{A}, \boldsymbol{Z})$$

$$\boldsymbol{\sigma_1} \perp\!\!\!\perp (\boldsymbol{B}, \boldsymbol{p}, \sigma, \boldsymbol{\sigma_0}, \hat{\boldsymbol{\Gamma}}) \mid (\boldsymbol{A}, \boldsymbol{Z})$$

Thus

$$p(\boldsymbol{\sigma_0}|\boldsymbol{A}, \boldsymbol{Z}) = p(\boldsymbol{\sigma_0}|\boldsymbol{A}, \boldsymbol{Z}, \boldsymbol{B}, \boldsymbol{p}, \sigma, \boldsymbol{\sigma_1}, \hat{\boldsymbol{\Gamma}})$$

$$p(\boldsymbol{\sigma_1}|\boldsymbol{A},\boldsymbol{Z}) = p(\boldsymbol{\sigma_1}|\boldsymbol{A},\boldsymbol{Z},\boldsymbol{B},\boldsymbol{p},\sigma,\boldsymbol{\sigma_0},\hat{\boldsymbol{\Gamma}})$$

Also for any positive integer $k \le K$, since $\sigma_{k0}$ is independent of $(\boldsymbol{A}_{k',*}, \boldsymbol{Z}_{k',*})$ for $k' \ne k$, we have

$$p(\sigma_{k0}|\boldsymbol{A},\boldsymbol{Z}) = p(\sigma_{k0}|\alpha_{k1},\ldots,\alpha_{kp},z_{k1},\ldots,z_{kp}).$$

Similarly

$$p(\sigma_{k1}|\boldsymbol{A},\boldsymbol{Z}) = p(\sigma_{k1}|\alpha_{k1},\ldots,\alpha_{kp},z_{k1},\ldots,z_{kp}).$$

Note

$$\alpha_{kj} \sim \begin{cases} N(0,\sigma_{k0}^2) & \text{if } z_{kj}=0, \\ N(0,\sigma_{k1}^2) & \text{if } z_{kj}=1 \end{cases},$$

with prior,

$$\sigma_{kq}^2 \sim \text{InvGamma}(\alpha_k^q, \beta_k^q),$$

where $q \in \{0,1\}$.

Then the posterior is,

$$\sigma_{kq}^2 \sim \text{InvGamma}\left(\alpha_k^q + \frac{1}{2}n_{kq},\ \beta_k^q + \frac{1}{2}\sum_{j=1}^{P}\alpha_{kj}^2\mathbb{1}\{z_{kj}=q\}\right),$$

where $n_{iq} := \sum_{j=1}^{P}\mathbb{1}\{z_{kj}=q\}$.

##### S1.4.4 Updating $p_k$

Note

$$\boldsymbol{p} \perp\!\!\!\perp (\boldsymbol{B},\sigma,\boldsymbol{\sigma_0},\boldsymbol{\sigma_1},\boldsymbol{A},\hat{\boldsymbol{\Gamma}}) \mid \boldsymbol{Z}$$

Therefore

$$p(\boldsymbol{p}|\boldsymbol{Z}) = p(\boldsymbol{p}|\boldsymbol{B},\sigma,\boldsymbol{\sigma_0},\boldsymbol{\sigma_1},\boldsymbol{A},\hat{\boldsymbol{\Gamma}},\boldsymbol{Z})$$

Since $p_k$ is independent of $\boldsymbol{Z}_{k',*}$ for $k' \ne k$, we have

$$p(p_k|\boldsymbol{Z}) = p(p_k|z_{k1},\ldots,z_{kp})$$

With prior

$$p_k \sim \text{Beta}(a_k, b_k),$$

and likelihood

$$z_{kj} \sim \text{Bernoulli}(p_k),$$

we have Posterior:

$$p_k \sim \text{Beta}(a_k + n_k,\ b_k + P - n_k),$$

where $n_k = \sum_{j=1}^{P} z_{kj}$.

##### S1.4.5 Updating $(\boldsymbol{A},\boldsymbol{Z})$

To sample $(\boldsymbol{A},\boldsymbol{Z})$ we first sample marginal of $\boldsymbol{Z}$ given everything but $\boldsymbol{A}$. Observe that

$$\begin{aligned} &p(\boldsymbol{Z}_{*,j},\hat{\boldsymbol{\Gamma}}|\boldsymbol{B},\boldsymbol{p},\sigma,\boldsymbol{\sigma_0},\boldsymbol{\sigma_1}) \\ =&p(\boldsymbol{Z}_{*,j},\hat{\boldsymbol{\Gamma}}_{*,j}|\boldsymbol{B},\boldsymbol{p},\sigma,\boldsymbol{\sigma_0},\boldsymbol{\sigma_1})\cdot\left(\prod_{j'\ne j}p(\hat{\boldsymbol{\Gamma}}_{*,j'}|\boldsymbol{B},\boldsymbol{p},\sigma,\boldsymbol{\sigma_0},\boldsymbol{\sigma_1}\right). \end{aligned}$$

Divide both sides by $p(\hat{\boldsymbol{\Gamma}})$. By independence among columns of $\hat{\boldsymbol{\Gamma}}$, we have

$$p(\boldsymbol{Z}_{*,j}|\hat{\boldsymbol{\Gamma}},\boldsymbol{B},\boldsymbol{p},\sigma,\boldsymbol{\sigma_0},\boldsymbol{\sigma_1}) = p(\boldsymbol{Z}_{*,j}|\hat{\boldsymbol{\Gamma}}_{*,j},\boldsymbol{B},\sigma,\boldsymbol{p},\boldsymbol{\sigma_0},\boldsymbol{\sigma_1}).$$

Therefore we can just inspect each $\boldsymbol{Z}_{*,j}$ on its own. We have a likelihood given by:

$$\begin{aligned}\hat{\boldsymbol{\Gamma}}_{*,j}|\boldsymbol{Z}_{*,j},\boldsymbol{\sigma_0},\boldsymbol{\sigma_1} &= (\boldsymbol{I}+\boldsymbol{B})\boldsymbol{A}_{*,j}+\boldsymbol{\epsilon}_{*,j}\\ &\sim \mathcal{N}(\boldsymbol{0},\ (\boldsymbol{I}+\boldsymbol{B})\boldsymbol{\Sigma}_{Z_{*,j}}(\boldsymbol{I}+\boldsymbol{B})^T+\boldsymbol{\Sigma}_j),\end{aligned} \tag{14}$$

where

$$\boldsymbol{\Sigma}_{Z_{*,j}} := \text{diag}(\sigma_{1z_{1j}},\dots,\sigma_{Kz_{Kj}}),$$

and we have prior

$$z_{kj} \overset{\text{iid}}{\sim} \text{Bernoulli}(p_k).$$

In cases where $K$ is small enough, we can easily compute the un-normalized probability weights for all possibilities of $Z_j$, and sample from a discrete distribution.

Now that we have a sample from the marginal of $\boldsymbol{Z}$ we can sample from the conditional of $\boldsymbol{A}$, which has prior of

$$\boldsymbol{A}_{*,j} \sim \mathcal{N}(\boldsymbol{0},\boldsymbol{\Sigma}_{Z_{*,j}}). \tag{15}$$

Recall that we have the linear Guassian model,

$$\hat{\boldsymbol{\Gamma}}_{*,j} = (\boldsymbol{I}+\boldsymbol{B})\boldsymbol{A}_{*,j}+\boldsymbol{\epsilon}_{*,j}$$

with noise distribution,

$$\boldsymbol{\epsilon}^T_{*,j} \sim \mathcal{N}(\boldsymbol{0},\boldsymbol{\Sigma}_j),$$

so we get a posterior similar to before of

$$\begin{aligned}\boldsymbol{A}_{*,j}|\hat{\boldsymbol{\Gamma}}_{*,j},\boldsymbol{B},\boldsymbol{\sigma}_0,\boldsymbol{\sigma}_1 &\sim \mathcal{N}(\boldsymbol{m},\boldsymbol{c}),\\ \boldsymbol{C} &= [(\boldsymbol{I}+\boldsymbol{B})^T\boldsymbol{\Sigma}_j^{-1}(\boldsymbol{I}+\boldsymbol{B})+\boldsymbol{\Sigma}^{-1}_{\boldsymbol{Z}_{*,j}}]^{-1},\\ \boldsymbol{m} &= \boldsymbol{C}(\boldsymbol{I}+\boldsymbol{B})^T\boldsymbol{\Sigma}_j^{-1}\hat{\boldsymbol{\Gamma}}_{*,j}.\end{aligned}$$

#### S1.5 Estimation for hyper-parameters

In our empirical studies, we find that the posterior distributions of $B$ can be sensitive to the choice of the hyper-parameters $(\alpha_k^0,\beta_k^0),(\alpha_k^1,\beta_k^1)$ and $(a_k,b_k)$. Here we take an empirical based approach to choose the hyper-parameters. Specifically, these hyperparameters will be set to the corresponding maximum likelihood estimators in the marginalized model, which can be obtained by the expectation-maximization (EM) algorithm. Recall that our model assumes for $k\in\{1,2,\dots,K\}$,

$$\alpha_{kj} \sim (1-p_k)\mathcal{N}(0,\sigma_{k0}^2)+p_k\mathcal{N}(0,\sigma_{k1}^2) \tag{16}$$

In particular,

$$\hat{\gamma}_{1j} = \alpha_{1j}+\epsilon_{1j} \sim (1-p_1)N(0,\sigma_{10}^2+\delta_{1j}^2)+p_1\mathcal{N}(0,\sigma_{11}^2+\delta_{1j}^2) \tag{17}$$

We first estimate the hyper-parameters for the outcome pleiotropies. In the E step, we compute the posterior distribution of the latent variables given our current estimates of parameters. In the M step we solve the maximization problem of the expectation of the log likelihood over the latent variables. To get rid of the identifiability issue, we initialize $\sigma_{10}^2$ and $\sigma_{11}^2$ to be far from each other. When solving for $\sigma_{10}^2$ and $\sigma_{11}^2$ in the M-step, we put the restriction that they can be at most 10 times larger than the their values from last iteration. Once we have obtained the point estimates of $(p_1,\sigma_{10},\sigma_{11})$, we choose the hyper-parameters for their priors such that the prior means are equal to the points estimates.

For $k\in\{2,3,\dots,K\}$, however, this is theoretically unachievable because $\hat{\alpha}_{kj}\equiv\alpha_{kj}+\epsilon_{kj}$ are no longer accessible. In practice we set $\hat{\alpha}_{kj}$ to $\hat{\gamma}_{kj}$ to determine the hyper-parameters. The priors will be biased especially when the genetic effects of a trait is largely mediated by an upstream trait that is already in the model. Nonetheless, our numerical simulations suggest that this bias may be small in most reasonable scenarios. We point out that one possible alternative approach is to use this as a first degree approximation and then use the Gibbs sampler to estimate $\boldsymbol{B}$ and then use that to estimate the direct genetic association, and repeat the expectation maximization. In the optimal settings, if the estimates of $\boldsymbol{B}$ in the first stage is close to the true value, this two-stage procedure will improve the precision of the algorithm. Indeed it turns out that this second stage estimation can usually be quite helpful in univariate Mendelian Randomization. On the other hand, the step of estimating the direct genetic association aggregates the noise from multiple exposures and makes it harder for EM algorithm to find meaning and reliable estimates for the hyperparameters. Therefore the two-state procedure is only recommended when the number of traits are small and the noise levels are fair.

##### S1.5.1 EM details

Recall that our model assumes for $k \in \{1, 2, \ldots, K\}$,

$$\alpha_{kj} \sim (1 - p_k)\mathcal{N}(0, \sigma_{k0}^2) + p_k\mathcal{N}(0, \sigma_{k1}^2)$$

In particular,

$$\hat{\gamma}_{1j} = \alpha_{1j} + \epsilon_{1j} \sim (1 - p_1)N(0, \sigma_{10}^2 + \delta_{1j}^2) + p_1\mathcal{N}(0, \sigma_{11}^2 + \delta_{1j}^2)$$

For simplicity, we will drop the subscript 1 for the time being.

**E-step.** Denote $\theta^{(t)} = (p^{(t)}, \sigma_1^{(t)}, \sigma_0^{(t)})$ .With latent variable $z_j \sim \text{Bernoulli}(p)$, where $\hat{\gamma}_j$ comes from the first distribution when $z_j = 1$, the membership probabilities are

$$w_j \equiv \mathbb{P}(z_j = 1|\theta^{(t)}) = \frac{p^{(t)}\phi(\hat{\gamma}_j; 0, (\sigma_1^{(t)})^2 + \delta_j^2)}{p^{(t)}\phi(\hat{\gamma}_j; 0, (\sigma_1^{(t)})^2 + \delta_j^2) + (1 - p^{(t)})\phi(\hat{\gamma}_j; 0, (\sigma_0^{(t)})^2 + \delta_j^2)},$$

where $\phi(; \mu, \sigma)$ is the density of normal distribution with mean $\mu$ and standard deviation $\sigma$. Then we can compute the expectation of the data-log-likelihood with respect to the distribution of $\boldsymbol{Z}$ given the data and $\theta^{(t)}$:

$$\begin{aligned}\mathbb{E}_{\boldsymbol{Z}}\left[\log L(\theta)|\hat{\gamma}_j, \theta^{(t)}\right] &= \sum_{j=1}^{P}\Big(w_j \log\left[p\phi(\hat{\gamma}_j; 0, \sigma_1^2 + \delta_j^2)\right] \\ &\qquad + (1 - w_j)\log\left[(1 - p)\phi(\hat{\gamma}_j; 0, \sigma_0^2 + \delta_j^2)\right]\Big) \\ &= \sum_{j=1}^{P}\Bigg(w_j\left[\log(p) - \frac{1}{2}\log(\delta_j^2 + \sigma_1^2) - \frac{\hat{\gamma}_j^2}{2(\delta_j^2 + \sigma_1^2)}\right] \\ &\qquad + (1 - w_j)\left[\log(1 - p) - \frac{1}{2}\log(\delta_j^2 + \sigma_0^2) - \frac{\hat{\gamma}_j^2}{2(\delta_j^2 + \sigma_0^2)}\right]\Bigg) + C,\end{aligned}$$

where $C$ is a constant that does not depend on $\theta$.

**M-step.** Taking derivative with respect to $p$ and setting it to zero yields

$$\sum_{j=1}^{P}\left(\frac{w_j}{p} - \frac{(1 - w_j)}{1 - p}\right) = 0 \implies p^{(t+1)} = \frac{\sum_{j=1}^{P} w_j}{P}$$

For $\sigma_1^2$, it is equivalent to minimize

$$\sum_{j=1}^{P}\left(\log(\delta_j^2+\sigma_1^2)+\frac{\hat{\gamma}_j^2}{(\delta_j^2+\sigma_1^2)}\right),$$

and similarly for $\sigma_0^2$ .The objectives can be maximized with a numerical optimization method.

#### S1.6 Extension to multiple traits at each time point

In this section we propose an extension of the Bayesian framework which allows multiple traits at a single time stage. Specifically, suppose there are $K$ time stages and $N$ traits in total. At each time stage $k$, there are $n_k$ exposures/outcomes of interests $X_{k1},\dots,X_{kn_k}$, with no internal interactions. Define $\boldsymbol{X}_1,\dots,\boldsymbol{X}_K$ as

$$\boldsymbol{X}_1=\begin{pmatrix}X_{11}\\X_{12}\\\vdots\\X_{1n_1}\end{pmatrix},\boldsymbol{X}_2=\begin{pmatrix}X_{21}\\X_{22}\\\vdots\\X_{2n_2}\end{pmatrix},\boldsymbol{X}_3=\begin{pmatrix}X_{31}\\X_{32}\\\vdots\\X_{3n_3}\end{pmatrix},\cdots,\boldsymbol{X}_K=\begin{pmatrix}X_{K1}\\X_{K2}\\\vdots\\X_{Kn_K}\end{pmatrix}.$$

Let

$$\gamma_{ki,j}=\text{argmin}_{\gamma}\text{Var}\left[X_{ki}-\gamma Z_j\right]$$

be the marginal association between $X_{ki}$ and $Z_j$. Define $\boldsymbol{\gamma}_{1,j},\dots,\boldsymbol{\gamma}_{K,j}$ as

$$\boldsymbol{\gamma}_{1,j}=\begin{pmatrix}\gamma_{11,j}\\\gamma_{12,j}\\\vdots\\\gamma_{1n_1,j}\end{pmatrix},\boldsymbol{\gamma}_{2,j}=\begin{pmatrix}\gamma_{21,j}\\\gamma_{22,j}\\\vdots\\\gamma_{2n_2,j}\end{pmatrix},\boldsymbol{\gamma}_{3,j}=\begin{pmatrix}\gamma_{31,j}\\\gamma_{32,j}\\\vdots\\\gamma_{3n_3,j}\end{pmatrix},\cdots,\boldsymbol{\gamma}_{K,j}=\begin{pmatrix}\gamma_{K1,j}\\\gamma_{K2,j}\\\vdots\\\gamma_{Kn_K,j}\end{pmatrix}$$

Similarly define $\boldsymbol{\alpha}_{1,j},\dots,\boldsymbol{\alpha}_{K,j}$ as

$$\boldsymbol{\alpha}_{1,j}=\begin{pmatrix}\alpha_{11,j}\\\alpha_{12,j}\\\vdots\\\alpha_{1n_1,j}\end{pmatrix},\boldsymbol{\alpha}_{2,j}=\begin{pmatrix}\alpha_{21,j}\\\alpha_{22,j}\\\vdots\\\alpha_{2n_2,j}\end{pmatrix},\boldsymbol{\alpha}_{3,j}=\begin{pmatrix}\alpha_{31,j}\\\alpha_{32,j}\\\vdots\\\alpha_{3n_3,j}\end{pmatrix},\cdots,\boldsymbol{\alpha}_{K,j}=\begin{pmatrix}\alpha_{K1,j}\\\alpha_{K2,j}\\\vdots\\\alpha_{Kn_K,j}\end{pmatrix}$$

Let

$$\boldsymbol{\Gamma}:=\begin{bmatrix}\boldsymbol{\gamma}_{1,1}&\boldsymbol{\gamma}_{1,2}&\cdots&\boldsymbol{\gamma}_{1,P}\\\vdots&\vdots&\vdots&\vdots\\\boldsymbol{\gamma}_{K,1}&\boldsymbol{\gamma}_{K,2}&\cdots&\boldsymbol{\gamma}_{K,P}\end{bmatrix}\in\mathbb{R}^{N\times P}$$

$$\boldsymbol{A}:=\begin{bmatrix}\boldsymbol{\alpha}_{1,1}&\boldsymbol{\alpha}_{1,2}&\cdots&\boldsymbol{\alpha}_{1,P}\\\vdots&\vdots&\vdots&\vdots\\\boldsymbol{\alpha}_{K,1}&\boldsymbol{\alpha}_{K,2}&\cdots&\boldsymbol{\alpha}_{K,P}\end{bmatrix}\in\mathbb{R}^{N\times P}$$

Similar to the previous setting, from the individual level equations we have

$$\boldsymbol{\Gamma}=\tilde{\boldsymbol{B}}\boldsymbol{\Gamma}+\boldsymbol{A},\tag{18}$$

where $\tilde{\boldsymbol{B}}$ is the matrix of coefficients. Furthermore, since there is no interactions between $X_{ki}$'s within each time stage $k$, we the coefficient matrix can be written as

$$\tilde{\boldsymbol{B}} = \begin{bmatrix} \mathbf{0}_{\boldsymbol{n_1}} & \mathbf{0} & \mathbf{0} & \mathbf{0} & \dots & \mathbf{0} \\ \tilde{\boldsymbol{B}}_2 & \mathbf{0}_{\boldsymbol{n_2}} & \mathbf{0} & \mathbf{0} & \dots & \mathbf{0} \\ & \tilde{\boldsymbol{B}}_3 & \mathbf{0}_{\boldsymbol{n_3}} & \mathbf{0} & \dots & \mathbf{0} \\ & & \tilde{\boldsymbol{B}}_4 & \mathbf{0}_{\boldsymbol{n_4}} & \dots & \mathbf{0} \\ & & \dots & & \ddots & \mathbf{0} \\ & & & \tilde{\boldsymbol{B}}_K & & \mathbf{0}_{\boldsymbol{n_K}} \end{bmatrix},$$

where $\mathbf{0}_{\boldsymbol{n_k}} \in \mathbb{R}^{n_k \times n_k}$ represents the square matrix of zeros, $\tilde{\boldsymbol{B}}_k \in \mathbb{R}^{n_k \times (\sum_i^k n_i)}$ represents the matrix of marginal associations between the $\boldsymbol{X}_k$ and $(\boldsymbol{X}_l)_{1 \le l < k}$.

Equivalently by defining $\boldsymbol{B} = (\boldsymbol{I} - \tilde{\boldsymbol{B}})^{-1} - \boldsymbol{I}$ we can write

$$\begin{aligned} \boldsymbol{\Gamma} &= (\boldsymbol{I} + \boldsymbol{B})\boldsymbol{A} && (19) \\ \hat{\boldsymbol{\Gamma}} &= (\boldsymbol{I} + \boldsymbol{B})\boldsymbol{A} + \boldsymbol{\epsilon}, && (20) \end{aligned}$$

where $\boldsymbol{B}$ is of the form

$$\boldsymbol{B} = \begin{bmatrix} \mathbf{0}_{\boldsymbol{n_1}} & \mathbf{0} & \mathbf{0} & \mathbf{0} & \dots & \mathbf{0} \\ \boldsymbol{B}_2 & \mathbf{0}_{\boldsymbol{n_2}} & \mathbf{0} & \mathbf{0} & \dots & \mathbf{0} \\ & \boldsymbol{B}_3 & \mathbf{0}_{\boldsymbol{n_3}} & \mathbf{0} & \dots & \mathbf{0} \\ & & \boldsymbol{B}_4 & \mathbf{0}_{\boldsymbol{n_4}} & \dots & \mathbf{0} \\ & & \dots & & \ddots & \mathbf{0} \\ & & & \boldsymbol{B}_K & & \mathbf{0}_{\boldsymbol{n_K}} \end{bmatrix}.$$

This is a direct conclusion from the fact that

$$\left(\boldsymbol{I} - \tilde{\boldsymbol{B}}\right)^{-1} - \boldsymbol{I} = \boldsymbol{I} + \sum_{k=1}^{\infty} \tilde{\boldsymbol{B}}^k - \boldsymbol{I} = \sum_{k=1}^{\infty} \tilde{\boldsymbol{B}}^k.$$

##### S1.6.1 Adaption of Gibbs Sampler

The implementation of the Gibbs sampler for the multivariate version is identical to the previous version except for the updates of $\boldsymbol{B}$, since we impose addition structure on $\boldsymbol{B}$. We make the following definitions:

- Define $N_k = \sum_{i=1}^k n_i$ for $k = 1, 2, \dots, K$.
- For any $i \in \{n_1 + 1, n_1 + 2, \dots, N_k\}$, define $f(i)$ to be the unique integer $k$ in $\{2, \dots, K\}$ such that $N_{k-1} < i \le N_k$. Define $g(i) = N_{f(i)}$.
- For any matrix $\boldsymbol{B} \in \mathbb{R}^{N \times N}$, given $n_1, \dots, n_k$, define

$$\text{BLvec}(\boldsymbol{B}) := \begin{pmatrix} B_{N_1+1,1} \\ B_{N_1+1,2} \\ \vdots \\ B_{N_1+1,g(N_1+1)} \\ B_{N_1+2,1} \\ B_{N_1+2,2} \\ \vdots \\ B_{N_1+2,g(N_1+2)} \\ \vdots \\ B_{N,1} \\ B_{N,2} \\ \vdots \\ B_{N,g(N)} \end{pmatrix}$$

which is the row-wise vectorization of matrix $\boldsymbol{B}$, excluding the deterministic zero values.

- For any $\boldsymbol{v} \in \mathbb{R}^N$, define

$$\text{BLstack}(\boldsymbol{v}) = \begin{pmatrix} \mathbf{0}_{n_1}^T & v_1 & 0 & \cdots & 0 \\ \mathbf{0}_{n_1}^T & v_2 & 0 & \cdots & 0 \\ \vdots & \vdots & \vdots & \vdots & 0 \\ \mathbf{0}_{n_1}^T & v_{g(N_1+1)} & 0 & \cdots & 0 \\ \mathbf{0}_{n_1}^T & 0 & v_1 & \cdots & 0 \\ \mathbf{0}_{n_1}^T & 0 & v_2 & \cdots & 0 \\ \vdots & \vdots & \vdots & \vdots & \vdots \\ \mathbf{0}_{n_1}^T & 0 & v_{g(N_1+2)} & \cdots & 0 \\ \vdots & \vdots & \vdots & \vdots & \vdots \\ \mathbf{0}_{n_1}^T & 0 & 0 & \cdots & v_1 \\ \vdots & \vdots & \vdots & \vdots & \vdots \\ \mathbf{0}_{n_1}^T & 0 & 0 & \cdots & v_{g(N)} \end{pmatrix}$$

Hence, we have

$$\boldsymbol{B}\boldsymbol{v} = [\text{BLvec}(\boldsymbol{B})^T \text{BLstack}(\boldsymbol{v})]^T$$

Then we show how to update $\boldsymbol{B}$. With prior

$$\text{BLvec}(\boldsymbol{B}) \sim \mathcal{N}(\mathbf{0}, \sigma^2 \boldsymbol{I}),$$

likelihood

$$\boldsymbol{\epsilon}_{*,j} \sim \mathcal{N}(\mathbf{0}, \boldsymbol{\Sigma}_j),$$

and

$$\text{vec}(\boldsymbol{\epsilon}) \sim \mathcal{N}\left(0, \begin{pmatrix} \boldsymbol{\Sigma}_1 & \mathbf{0} & \cdots & \mathbf{0} \\ \mathbf{0} & \boldsymbol{\Sigma}_2 & \cdots & \mathbf{0} \\ \vdots & & \ddots & \vdots \\ \mathbf{0} & \mathbf{0} & \cdots & \boldsymbol{\Sigma}_P \end{pmatrix}\right),$$

we have

$$\begin{aligned} \hat{\boldsymbol{\Gamma}} &= (\boldsymbol{I} + \boldsymbol{B})\boldsymbol{A} + \boldsymbol{\epsilon} \\ \hat{\boldsymbol{\Gamma}} - \boldsymbol{A} &= \boldsymbol{B}\boldsymbol{A} + \boldsymbol{\epsilon} \\ \text{vec}(\hat{\boldsymbol{\Gamma}} - \boldsymbol{A})^T &= \text{BLvec}(\boldsymbol{B})^T \tilde{\boldsymbol{A}} + \text{vec}(\boldsymbol{\epsilon})^T \\ \text{vec}(\hat{\boldsymbol{\Gamma}} - \boldsymbol{A}) &= \tilde{\boldsymbol{A}}^T \text{BLvec}(\boldsymbol{B}) + \text{vec}(\boldsymbol{\epsilon}), \end{aligned}$$

where

$$\tilde{\boldsymbol{A}} = \begin{pmatrix} \text{BLstack}(\boldsymbol{A}_{*,1}) & \text{BLstack}(\boldsymbol{A}_{*,2}) & \cdots & \text{BLstack}(\boldsymbol{A}_{*,P}) \end{pmatrix}$$

So

$$\begin{aligned} \text{BLvec}(\boldsymbol{B}) &\sim \mathcal{N}(\boldsymbol{m}, \boldsymbol{C}) \\ \boldsymbol{C} &= (\tilde{\boldsymbol{A}}\boldsymbol{\Sigma}^{-1}\tilde{\boldsymbol{A}}^T + \sigma^{-2}\boldsymbol{I})^{-1} \\ \boldsymbol{m} &= \boldsymbol{c}\tilde{\boldsymbol{A}}\boldsymbol{\Sigma}^{-1}\text{vec}(\hat{\boldsymbol{\Gamma}} - \boldsymbol{A}) \end{aligned}$$

The other parts are identical to the previous case.

### S2 Additional details of simulation and real data analysis

#### S2.1 Simulation setup

Our simulation is based on multiple real GWAS summary statistics datasets. For the $K = 3$ scenario, we select SNPs using data from GIANT Adult BMI and EGG childhood BMI GWAS datasets [6]. The chosen SNPs are then associated with traits including 8-year-old BMI from MoBa, adult BMI from UK Biobank, and Type II diabetes (T2D) from DIAGRAM to generate "true" direct associations with the exposure and outcome traits. Extending to $K = 4$, the same SNPs and traits are retained, with an additional inclusion of the 1-year-old BMI GWAS trait from MoBa as the exposure trait. In the simulation scenario involving multivariate traits at each time point, for SNP selection we use summary statistics from low-density lipoprotein cholesterol (LDL-C), high-density lipoprotein cholesterol (HDL-C), and triglycerides in Genetic Epidemiology Research on Adult Health and Aging (GERA) [7]. The simulation comprises three time points: a first stage with childhood traits with LDL-C, HDL-C, and TG summary statistics from [8]; a second stage with adulthood traits with LDL-C, HDL-C, and TG summary statistics from [9]; and a third stage with the stroke trait as the outcome, utilizing summary statistics from [10].

To generate GWAS summary data for simulation, we first select SNPs based on the selection files with LD clumping [11] and a p-value cutoff at 0.01. For each selected SNP $j$ and each exposure $k$, we record the estimated effects $\hat{\gamma}_{kj}^{real}$ along with their corresponding standard errors $\hat{\delta}_{kj}$ in the exposure and outcome files. Subsequently, we set "true" direct associations $\alpha_{kj} = \text{sign}(\hat{\gamma}_{kj}^{real})(|\hat{\gamma}_{kj}^{real}| - 0.01)_+$, and we perform a random shuffle of $\alpha_{kj}$ across $j$ within each trait $k$ to ensure independence across traits. Once the matrix $\boldsymbol{A}$ is generated, we create $\boldsymbol{\Gamma}$ following our linear model with a pre-specified matrix $\boldsymbol{B}$. In the simulation, the matrix $\boldsymbol{B}$ is typically designed as an approximation of the true relationships among the traits in the real data. Finally, we simulate our synthetic summary statistics $\hat{\gamma}_{kj}^{simu} \sim \mathcal{N}(\gamma_{kj}, \hat{\delta}_{kj}^2)$ independently across all SNPs and traits. The results of experiments are based on 100 replications.

At each replication, in order to mimic the real case, we would like to run the algorithms with p-values cutoffs for the selection of SNPs. Unlike the real cases, where independent selection files are usually utilized to compute the instrumental strength without selection bias, we directly set p-values for each SNP by computing p-values from $\gamma_{kj}/\hat{\delta}_{kj}$ and taking the minimum of p-values across $k$. This can be heuristically considered as if we have independent selection files with no measurement errors. Specially, the p-values for each SNP $j$ is computed by

$$\min_{k<K} \left(2 \cdot \Phi^{-1}(-|\gamma_{kj}/\hat{\delta}_{kj}|)\right),$$

where $\Phi$ is the Gaussian cumulative distribution function. Bonferroni correction is then applied to select the significant SNPs under different p-values cutoffs.

#### S2.2 Sensitivity analysis to model assumptions

An important assumption of FLOW-MR is the independence of direct genetic effects across all traits. However, as discussed in earlier works on correlated pleiotropy [REF], such an independence assumption can be violated if there are hidden confounding pathways affecting multiple traits.

We perform an additional simulation to assess the robustness of FLOW-MR to the violation of the independence assumption. We consider a setting with $K = 3$ where direct genetic effects on the earliest two traits ($\alpha_{1j}$, $\alpha_{2j}$) are substantially correlated. Specifically, when we generate the synthetic data by soft-thresholding real GWAS summary statistics following Section S2.1 and the true causal relationship in Figure 3a of the main text, we only shuffle $\alpha_{3j}$ across $j$ so that it is independent from $\alpha_{1j}$ and $\alpha_{2j}$, while $\alpha_{1j}$ and $\alpha_{2j}$ are still correlated with an empirical correlation $\text{Corr}(\alpha_{1j}, \alpha_{2j}) = 0.45$.

When $X_2$ is the outcome, we observe that none of the competing methods, including those designed to handle correlated pleiotropy (e.g., MVMR-Horse, MVMR-cML), achieve the nominal 0.95% coverage (Figure S6a). Compared to other methods, FLOW-MR demonstrates higher coverage and lower estimation bias, indicating relative robustness to correlated pleiotropy.

When $Y = X_3$ is the outcome, where the pleiotropic effects on $Y$ are independent of the genetic associations on $X_1$ and $X_2$, FLOW-MR again matches GRAPPLE in maintaining the desired coverage—consistent with our main-text results (Figure S6b). These findings suggest that although FLOW-MR models the entire system jointly, its inference for a specific outcome remains robust to correlated direct genetic effects among other exposures, similar to MVMR methods.

#### S2.3 Computational cost of FLOW-MR

FLOW-MR relies on Bayesian Gibbs sampling, which can be computationally intensive for large-scale data. To address this, we have implemented FLOW-MR with Rcpp and parallelized the Gibbs sampler across four MCMC chains running on separate CPU cores by default. This design keeps FLOW-MR computationally feasible; for example, in a demanding setting with seven traits and 600 SNPs, the entire procedure finishes in roughly 17 minutes.

To benchmark computational cost (Figure S2.1), we compared FLOW-MR with other MVMR methods in both the $K = 3$ and multivariate simulation scenarios. Our results show that FLOW-MR is faster than MVMR-Horse (another Bayesian approach) and MVMR-cML. Notably, the reported runtime for FLOW-MR encompasses the joint estimation of all effects, whereas the computational cost shown for the MVMR methods is only for the final layer, where $X_K$ is the outcome. These findings suggest that FLOW-MR is suitably efficient and scalable for large datasets.

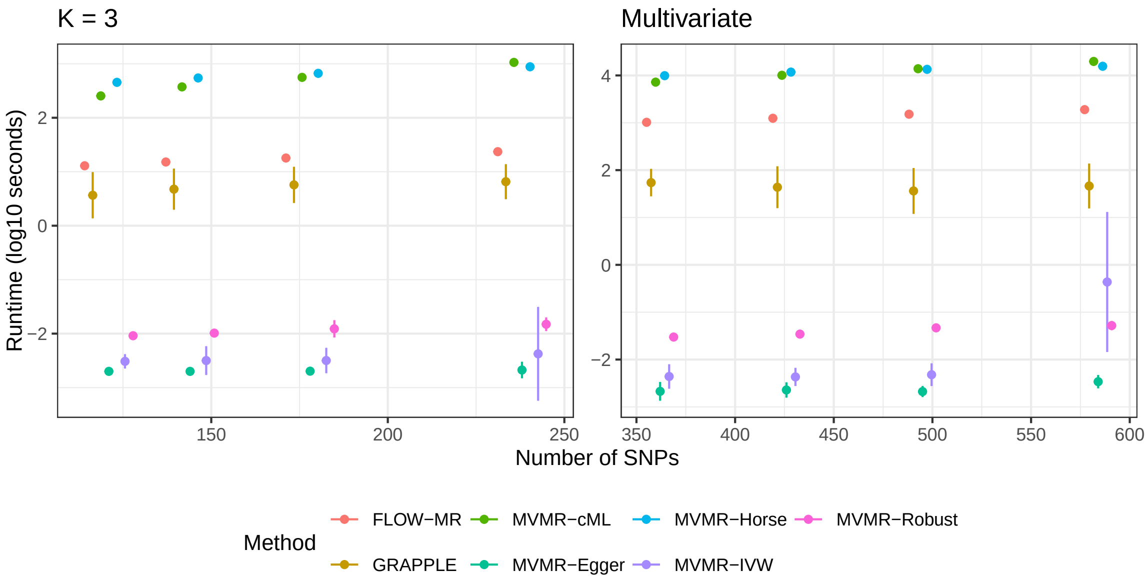


Figure S2.1: Compuational cost of different methods. FLOW-MR is run for 6,000 iterations, with the first 3,000 serving as a warm-up period. Error bars reflect two times standard derivations over 100 repeated experiments.

#### S2.4 Additional package information

The calculation of genetic correlation is carried out by the LDSC package developed in [12] and is available at (`https://github.com/bulik/ldsc`). All the plots in the paper are produced by `ggplot2` [13] (`https://ggplot2.tidyverse.org`) and `corrplot2` [14] (`https://github.com/taiyun/corrplot`, `https://github.com/mkanai/ldsc-corrplot-rg`).

#### Supplementary figures

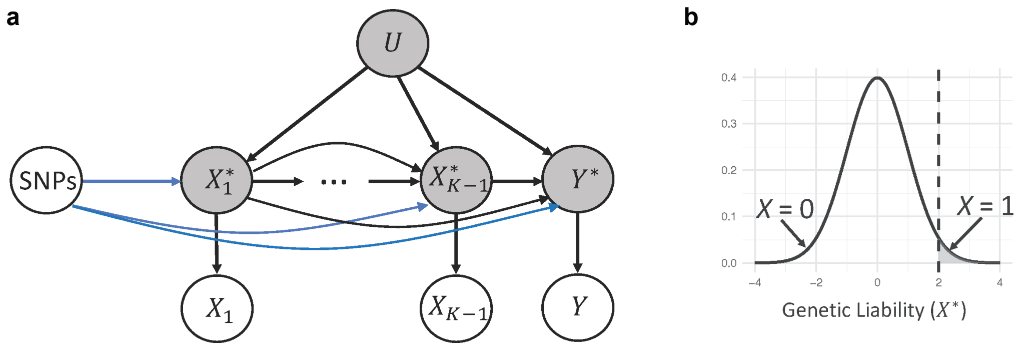


Figure S1: MR framework for binary data. a) Underlying causal DAG when some of the traits are binary. If $X_k$ is a continuous trait then the liability $X_k^\star = X_k$. If $X_k$ is a binary trait, then $X_k^\star$ is the underlying continuous liability. We assume that the causal relationships are between the liabilities. c) The liability threshold model. The binary trait $X_k = 1$ if and only if $X_k^\star > c$ for some threshold $c$.

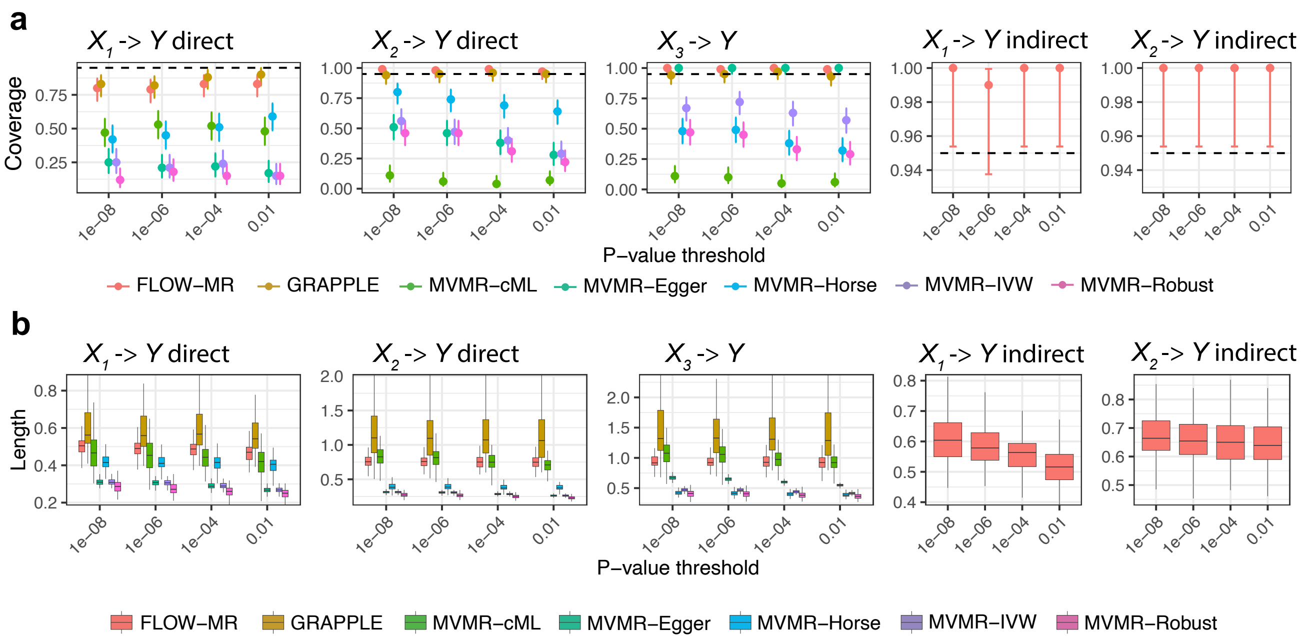


Figure S2: Simulation results for $K = 4$. a) Empirical coverage of 95% credible intervals of all direct/indirect effects of the exposures on $Y$ over 100 repeated simulations. The error bars are the 95% confidence intervals of the coverage. b) Boxplots of lengths of credible intervals over repeated simulations.

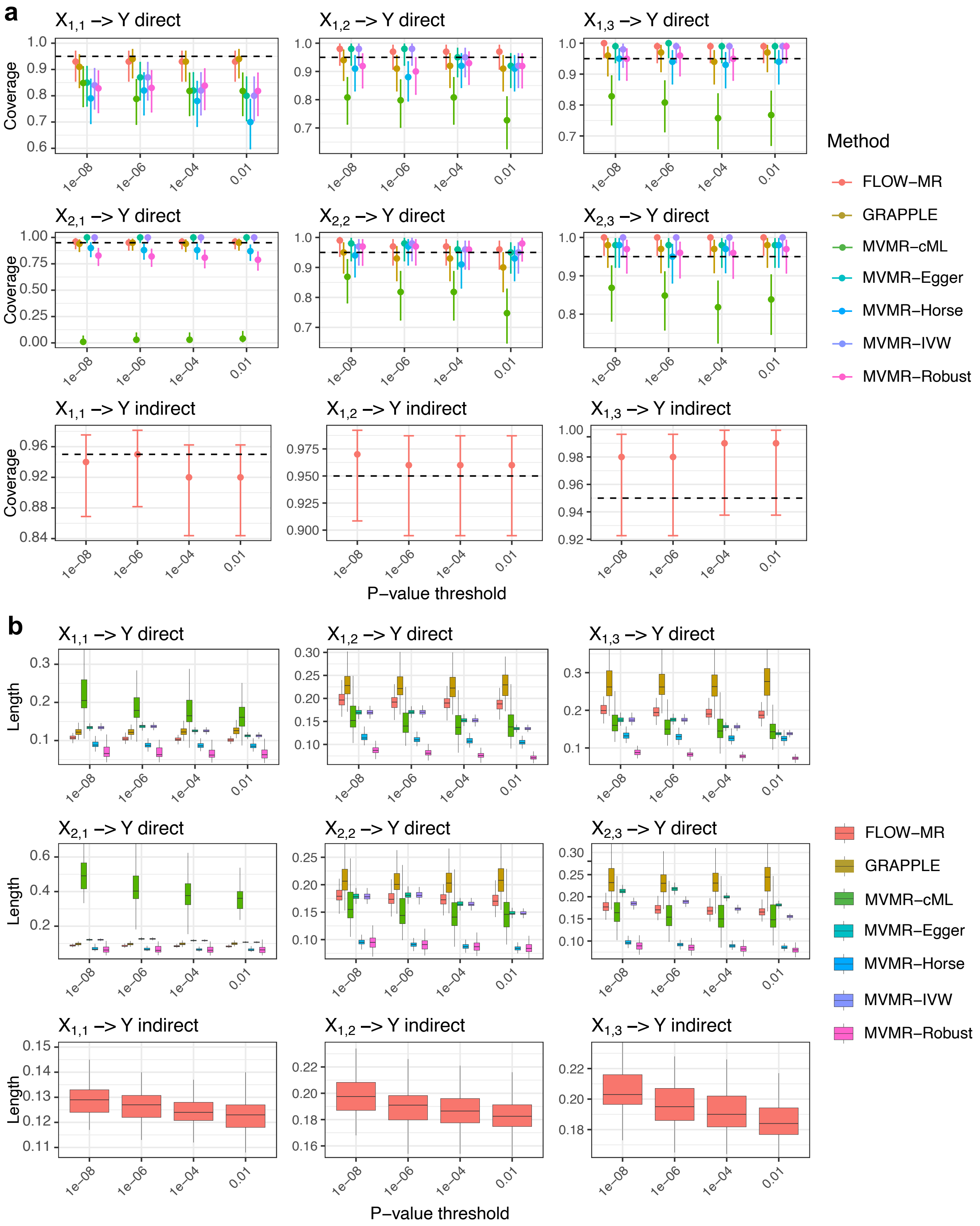


Figure S3: Simulation results for the multivariate case. a) Empirical coverage of 95% credible intervals of all direct/indirect effects of the exposures on $Y$ over 100 repeated simulations. The error bars are the 95% confidence intervals of the coverage. b) Boxplots of lengths of credible intervals over repeated simulations.

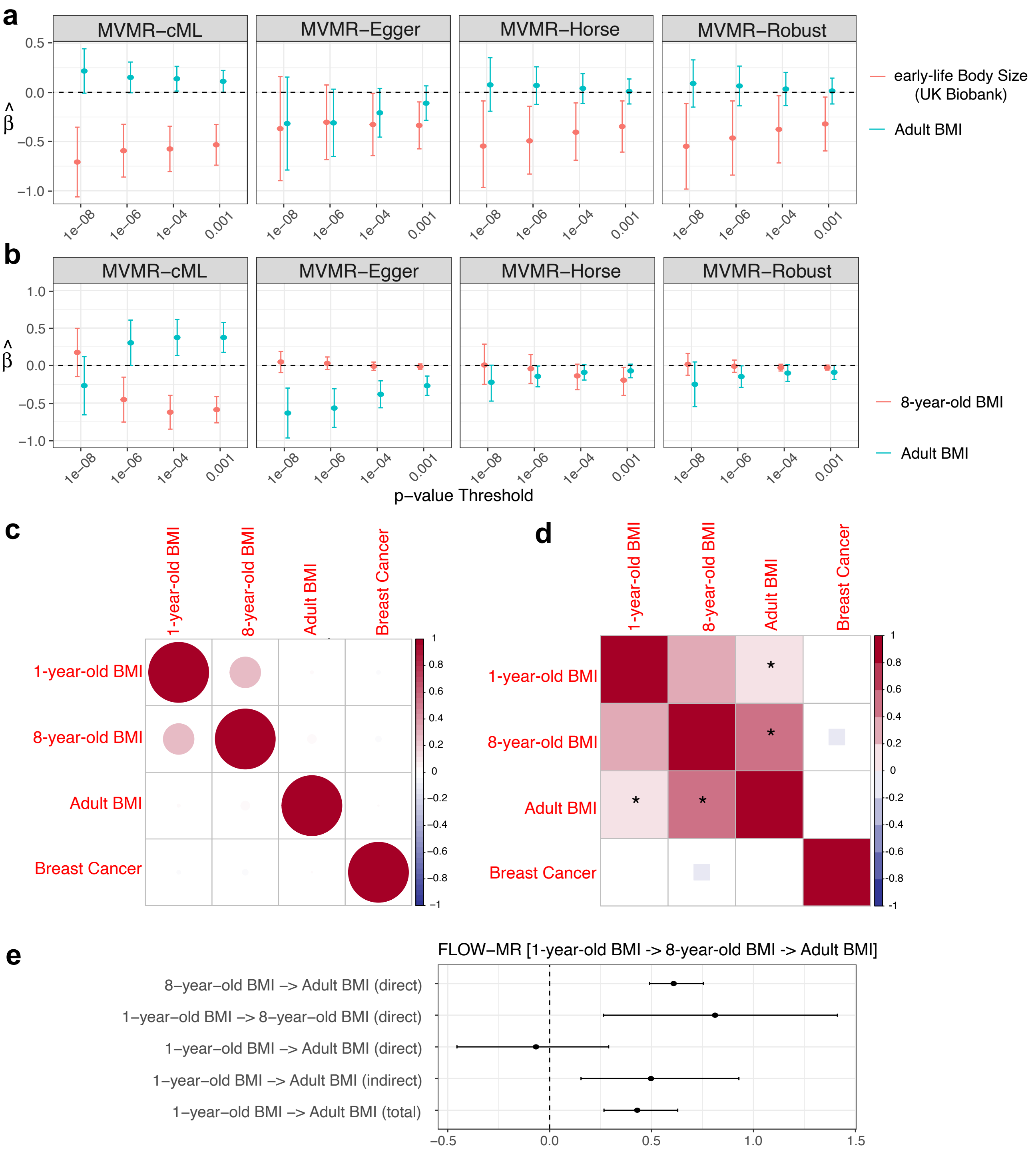


Figure S4: Additional results for the breast cancer case study. a) 95% Confidence / credible intervals of childhood body size and adult BMI on breast cancer risk estimated by 4 additional MVMR methods. b) 95% Confidence / credible intervals of childhood BMI and adult BMI on breast cancer risk estimated by 4 additional MVMR methods. c) Estimated noise correlation across traits. d) Estimated pairwise genetic correlations across traits. A '*' indicates significantly correlated pairs at significance level 0.05. e) Estimated noise correlation across traits. e) 95% credible intervals estimated by FLOW-MR for direct, indirect, and total effects among 1-year-old BMI, 8-year-old BMI, and adult BMI, with FLOW-MR applied to only these three traits.

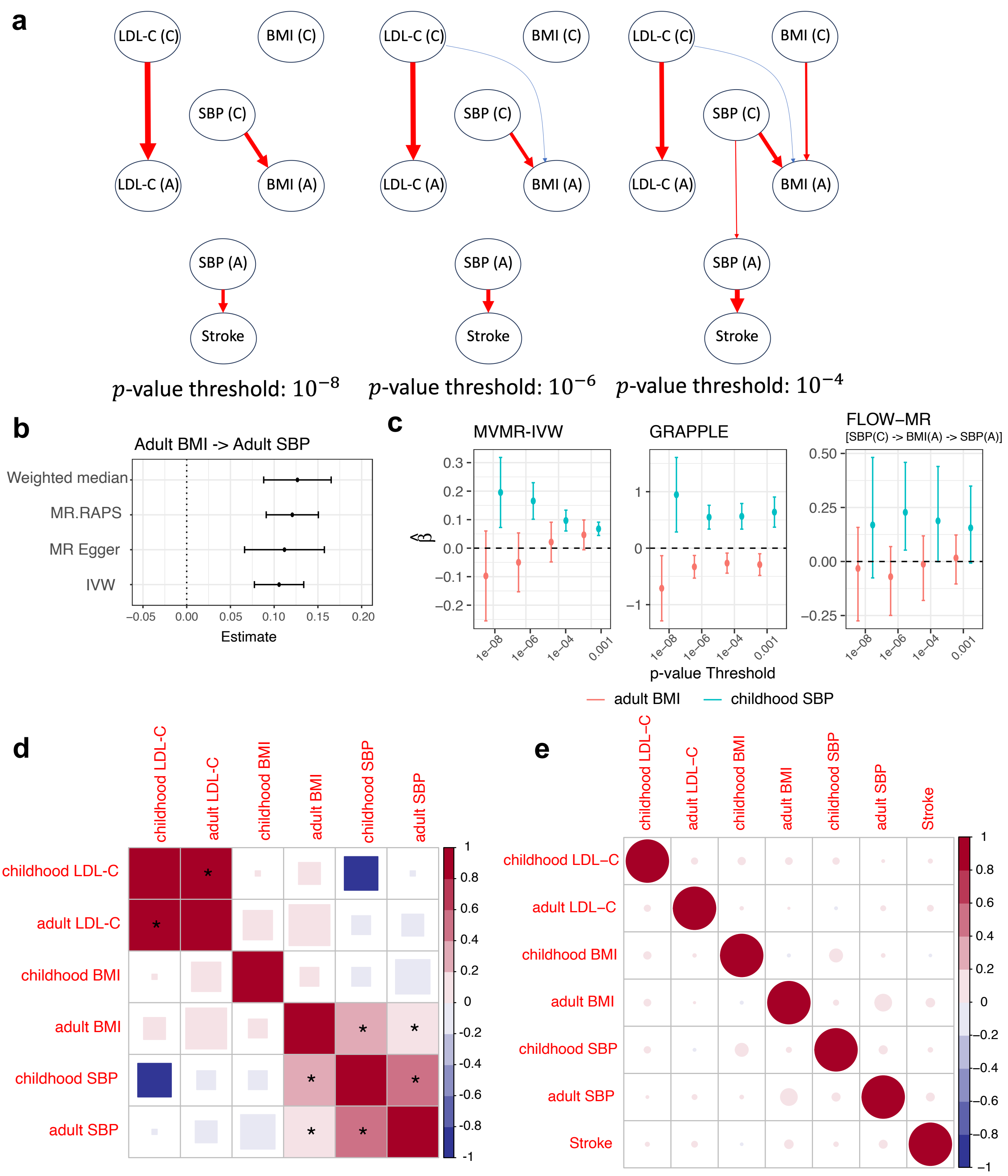


Figure S5: Additional results for the stroke case study. a) Estimated causal DAGs with different selection p-value thresholds. Arrows correspond to the significant effects, with thicker arrows indicating larger effects. Positive effects are shown in red, and negative effects in blue. b) Univariate MR 95% confidence intervals for the causal effect of adult BMI on adult SBP. c) 95% confidence intervals for the joint causal effects of adult BMI and childhood SBP on adult SBP using three methods. FLOW-MR is performed using only the three continuous traits involved. d) Estimated pairwise genetic correlations across traits. A '*' indicates significantly correlated pairs at significance level 0.05.

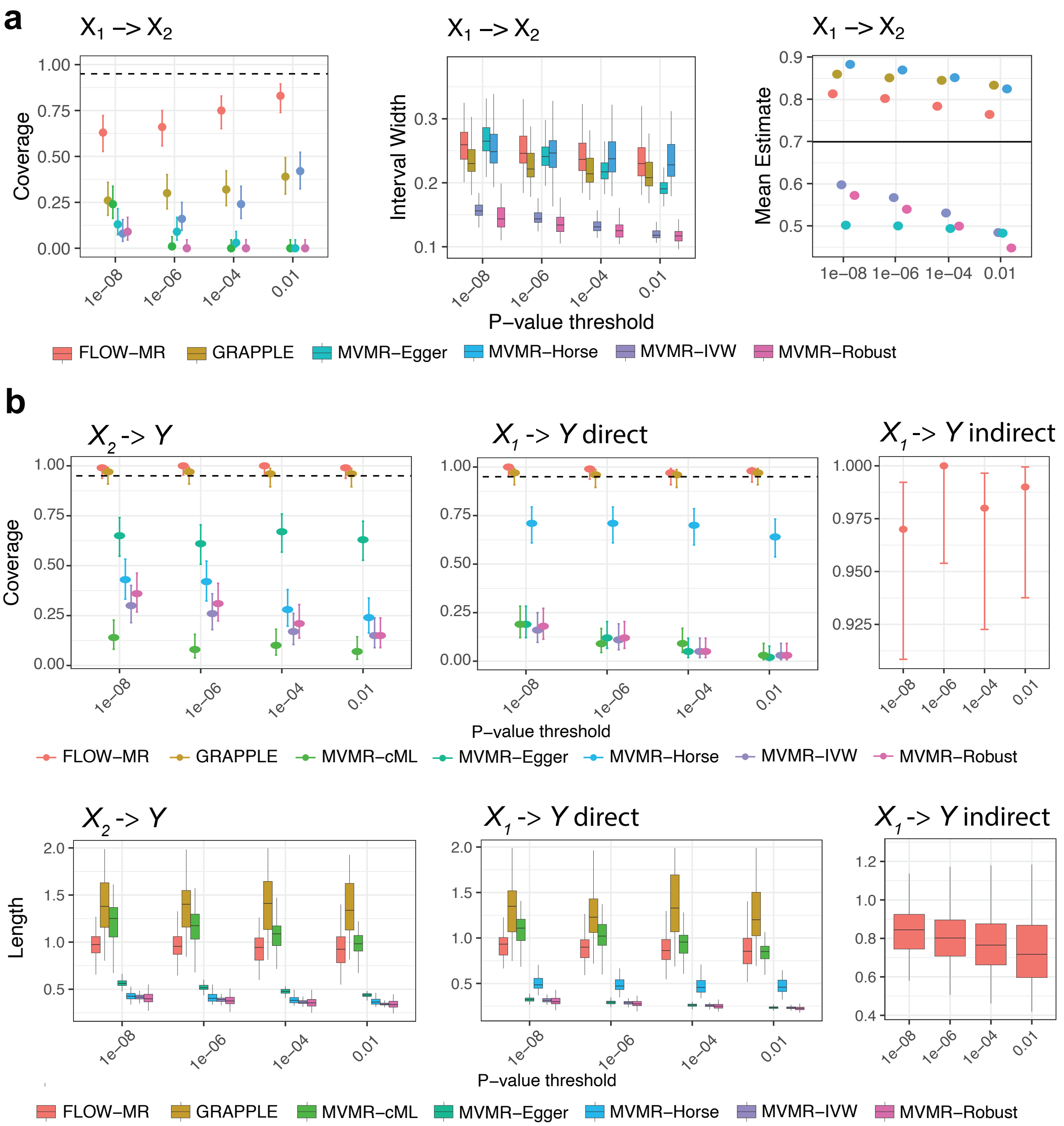


Figure S6: Simulation results for $K = 3$ under correlated direct genetic effects between $X_1$ and $X_2$. a) Empirical coverage and average 95% CI length, along with the mean estimated causal effect of $X_1$ on $X_2$ across different methods. b) Empirical coverage for the 95% CI of the direct causal effects of $X_1$ and $X_2$ on $Y = X_3$. The error bars indicate 95% confidence intervals for the true coverages. b) Boxplots of the 95% CI lengths for the direct causal effects of $X_1$ and $X_2$ on $Y = X_3$ across different methods.

#### Supplementary tables

Table S1: Number of SNPs used in simulations and real data analysis at any given selection p-value threshold

| Selection p-value thresholds | $10^{-8}$ | $10^{-6}$ | $10^{-4}$ | $10^{-2}$ |
|---|---|---|---|---|
| Simulation (K=3) | 121 | 144 | 178 | 238 |
| Simulation (K=4) | 158 | 174 | 205 | 236 |
| Simulation (Multivariate) | 362 | 426 | 495 | 584 |

| Selection p-value thresholds | $10^{-8}$ | $10^{-6}$ | $10^{-4}$ | $10^{-3}$ |
|---|---|---|---|---|
| Breast cancer (K=3, body size) | 56 | 119 | 305 | 570 |
| Breast cancer (K=3, BMI) | 56 | 118 | 304 | 571 |
| Breast cancer (K=4) | 55 | 117 | 303 | 570 |

| Selection p-value thresholds | $10^{-8}$ | $10^{-6}$ | $10^{-4}$ |
|---|---|---|---|
| Stroke | 122 | 210 | 341 |

Table S2: Conditional F statistics of each exposure in simulations and case studies.

| **Selection p-values** | $10^{-8}$ | $10^{-6}$ | $10^{-4}$ | $10^{-2}$ |
|---|---|---|---|---|
| **K = 3 (Simulation)** | | | | |
| $X_1$ | 1.86 | 1.76 | 1.55 | 1.60 |
| $X_2$ | 3.11 | 3.08 | 2.81 | 4.17 |
| **K = 4 (Simulation)** | | | | |
| $X_1$ | 2.17 | 2.16 | 2.01 | 1.96 |
| $X_2$ | 1.37 | 1.41 | 1.36 | 1.26 |
| $X_3$ | 2.18 | 2.34 | 2.49 | 2.47 |
| **Multivariate (Simulation)** | | | | |
| $X_{1,1}$ | 2.38 | 2.20 | 2.02 | 1.86 |
| $X_{1,2}$ | 1.58 | 1.55 | 1.43 | 1.40 |
| $X_{1,3}$ | 1.44 | 1.39 | 1.35 | 1.31 |
| $X_{2,1}$ | 3.78 | 3.74 | 3.50 | 3.36 |
| $X_{2,2}$ | 2.55 | 2.60 | 2.43 | 2.69 |
| $X_{2,3}$ | 2.63 | 2.57 | 2.62 | 2.72 |

| **Selection p-values** | $10^{-8}$ | $10^{-6}$ | $10^{-4}$ | $10^{-3}$ |
|---|---|---|---|---|
| **Breast Cancer (K = 3, body size)** | | | | |
| Early-life Body Size (UK Biobank) | 8.53 | 7.74 | 4.98 | 3.69 |
| Adult BMI | 8.86 | 8.83 | 5.79 | 4.32 |
| **Breast Cancer (K=3, BMI)** | | | | |
| 8-year-old BMI | 0.99 | 1.21 | 1.22 | 1.10 |
| Adult BMI | 1.50 | 3.17 | 5.36 | 7.03 |
| **Breast Cancer (K=4)** | | | | |
| 1-year-old BMI | 1.55 | 1.45 | 1.10 | 0.96 |
| 8-year-old BMI | 0.88 | 1.09 | 1.01 | 0.89 |
| Adult BMI | 1.50 | 3.61 | 6.01 | 7.61 |

| **Selection p-values** | $10^{-8}$ | $10^{-6}$ | $10^{-4}$ |
|---|---|---|---|
| **Stroke** | | | |
| LDL-C (C) | 1.35 | 1.25 | 1.35 |
| BMI (C) | 1.42 | 1.33 | 1.17 |
| SBP (C) | 1.03 | 1.12 | 1.08 |
| LDL-C (A) | 1.66 | 1.68 | 1.88 |
| BMI (A) | 4.55 | 5.81 | 7.13 |
| SBP (A) | 3.95 | 3.99 | 5.00 |
